# The danger zone: the joint trap of incomplete lineage sorting and long-branch attraction in resolving the Gondwanan origin of Rafflesiaceae and Apodanthaceae

**DOI:** 10.1101/2024.08.07.606681

**Authors:** Liming Cai, Natalia Pabón-Mora, Juan F. Alzate, Liang Liu, Charles C. Davis

## Abstract

Two key factors have been implicated as major impediments to phylogenomic inference: incomplete lineage sorting (ILS)—especially in cases where clades are in the so-called anomaly zone—and erroneous gene tree estimation—commonly manifested by long- branch attraction in the Felsenstein zone. Rafflesiaceae (Malpighiales) is an iconic parasitic plant clade whose species occur west of Wallace’s line in tropical Southeast Asia. This clade has been notoriously difficult to place phylogenetically and is nested within an explosive ancient radiation, rendering the Malpighiales one of the thorniest nodes across the angiosperm Tree of Life. The parasitic family Apodanthaceae has recently been positioned as sister to Rafflesiaceae, offering the hope to stabilize their placements. Here, using a dataset of 2,135 genes and complex species tree inference methods, the monophyletic Rafflesiaceae+Apodanthaceae clade is variously placed with Euphorbiaceae, Peraceae, Putranjivaceae, and Pandaceae. Such unstable placements appear to be the result of excessive levels of rate heterogeneity and ILS, which contribute to a phylogenetic “danger zone” where simulation suggests that current methods and genomic data will never provide a tidy phylogenetic resolution for species trees alike. Despite the topological uncertainty, however, our divergence time estimation identifies a mid-Cretaceous origin of stem group Rafflesiaceae and Apodanthaceae, not only making them the oldest parasitic plant lineage reported to date, but also suggests a likely Gondwana vicariance scenario to explain their disjunct distribution in South America, Africa, Australia, and Northern India.

## Introduction

Adaptive radiation is one of the most fascinating phenomena in evolutionary ecology; it occurs when a single lineage rapidly diverges into many descendant lineages each with substantial changes in their morphology, behavior, and physiology (Schluter 2000; Losos and Mahler 2010). Well-known examples include three-spined sticklebacks, Galapagos finches, Caribbean anoles, and Hawaiian silverswords and lobeliads (Schluter and McPhail 1992; Baldwin 1997; Grant and Grant 2007; Givnish et al. 2009; Losos 2011). Despite the intriguing nature of adaptive radiations, such clades often present challenges for phylogenetic inference because they frequently exhibit a combination of short and long branches resulting from extensive accumulation of mutations (Glor 2010; Walker and Smith 2023). Species trees involving adaptive radiations are likely to fall within the “anomaly zone” in which the most common gene tree differs from the species tree, resulting in a potentially high amount of incomplete lineage sorting (ILS) (Degnan and Rosenberg 2006). In addition, mutational saturation and rate heterogeneity among descendant branches in these radiations can lead to various systematic errors, including long-branch attraction and compositional bias (Kapli et al. 2020).

These factors are further confounded when the radiation is ancient, as in the case of Malpighiales—an apparent explosive Cretaceous radiation that has produced plants with diverse morphologies ranging from tropical mangroves (Rhizophoraceae) to submerged aquatics (Podostemaceae) (Davis et al. 2005; Wurdack and Davis 2009; Cai et al. 2020). Among the strangest of all malpighialean members is the parasitic family, Rafflesiaceae, whose species produce the world’s largest flowers (Davis et al. 2007; Nikolov and Davis 2017). Until recently, the large-flowered Rafflesiaceae sensu stricto (up to 1 m) were thought to be the only parasites within Malpighiales (Barkman et al. 2004; Davis and Wurdack 2004; Davis et al. 2007; Wurdack and Davis 2009), but Alzate et al. (2024) demonstrated the surprising placement of the small-flowered endoparasite Apodanthaceae species (less than 0.5 cm) within Malpighiales as sister to Rafflesiaceae. Their hypothesized synapomorphies include most notably the endophytic habit (endoparasite) where the plant persists as strands of cells with no distinguishable vegetative organs embedded in their hosts; until their flowers emerge briefly for sexual reproduction (Kuijt 1969; Nikolov et al. 2013; Thorogood et al. 2021; Teixeira-Costa and Davis 2021; González and Pabón-Mora 2017; Alzate et al. 2024). However, the two families have very distinct geographic distributions and host preferences. Rafflesiaceae parasitize vines of Tetrastigma (Vitaceae) and grow west of Wallace’s line in tropical Southeast Asia (Kuijt 1969; Nikolov et al. 2013). In contrast, Apodanthaceae exhibits a pantropical distribution with disjunct occurrences in subtropics and the Mediterranean. Members of Apodanthaceae parasitize Salicaceae and diverse Fabaceae shrubs and trees (Bellot and Renner 2014).

Reconstructing the early evolutionary history of these two parasitic families, including their phylogenetic placements and divergence time, will provide important insights into the biogeographical and ecological processes shaping these extraordinary parasites. Yet the path to their resolution has been slow. Rafflesiaceae has been proposed as sister to Passifloraceae and Euphorbiaceae sensu stricto within Malpighiales (Barkman et al. 2004; Nickrent et al. 2004; Davis et al. 2007; Bendiksby et al. 2010) and Apodanthaceae was a widely accepted member of Cucurbitales (Filipowicz and Renner 2010). The lack of genomic resources, excessive rate heterogeneity, and horizontal gene transfer have substantially impeded these previous investigations. The recently available genomic and transcriptomic resources in both families provide a promising opportunity to expand the toolkit for resolving their placement. In particular, the addition of Apodanthaceae offers hope of breaking up the long branch associated with stem group Rafflesiaceae. Despite this promise, there are four genomic aspects that will likely confound phylogenomic investigations. First, nearly half of the conserved genes in Rafflesiaceae (and likely in Apodanthaceae) have been lost due to the evolution of endoparasitism (Cai et al. 2021a), which greatly restricts the repertoire of genes for phylogenomic research. Moreover, these gene losses are overrepresented in housekeeping functions such as photosynthesis and plastid organization, which may lead to incorrect ortholog assessments during phylogenetic analyses (Xiong et al. 2022). Second, parasitic plants are known to acquire foreign genetic materials from their hosts via horizontal gene transfer (HGT) (Davis and Wurdack 2004; Xi et al. 2012; Xi et al. 2013). The incorrect placement of Apodanthaceae in Cucurbitales is likely an outcome of horizontally transferred matR and nuclear 18S from ancestral hosts (Filipowicz and Renner 2010). Third, the well-characterized high substitution rate in Rafflesiaceae may lead to spurious species tree reconstruction because of long-branch attraction (Nickrent and Starr 1994; Nickrent and Duff 1996). This phenomenon was shown in all three cellular genomes using representative genes across multiple independently evolved parasitic lineages (Young and dePamphilis 2005; Bromham et al. 2013) and has been demonstrated to span nearly every gene across the entire nuclear genome in Rafflesiaceae (Cai et al. 2021). Fourth, the unusual AT-rich (GC% = 24%) genome content of Rafflesiaceae likely contributes to compositional bias that may incorrectly group distantly related taxa as close relatives (Foster and Hickey 1999; Knight et al. 2001; Wang et al. 2004). Little is known regarding the substitution rates and AT content in Apodanthaceae. These confounding factors make the placement of this enigmatic clade incredibly challenging, but this is further compounded by the nature of their inclusion in the large and diverse order Malpighiales.

The ancient radiation of Malpighiales includes nine of the ten most phylogenetically recalcitrant angiosperm clades (Smith et al. 2013; Cai et al. 2019; Cai et al. 2020), easily placing its members as the most extraordinary and difficult phylogenomic problem to resolve in angiosperms. In addition, the potentially greatly accelerated substitution rates in Rafflesiaceae and Apodanthaceae combined with the previously summarized genomic anomalies attributable to their parasitic lifestyle make this an even greater intractable phylogenomic problem. This parasitic plant clade falls into a joint trap that we term “the danger zone” of ILS and long-branch attraction. The “danger zone” represents a perturbing combination not conducive to achieving phylogenetic resolution. But we envision that a careful assessment of these factors will yield insights into how to tackle an enormous phylogenomic challenge, which is likely to be found elsewhere in the tree of life. Here, using 2,135 genes including a broad sampling of Malpighiales, we tackle the joint problem of ILS and long-branch attraction using a combination of approaches that accommodate genealogical heterogeneity among loci and rate heterogeneity among sites. Our divergence time estimation demonstrated an ancient Gondwanan divergence of Rafflesiaceae and Apodanthaceae providing insight into their disjunct distribution and distinct host preference.

## RESULTS

### Phylogenomic datasets

By combining data from new RNA sequencing data from Apodanthaceae and two published genome and transcriptome studies within Malpighiales (Cai et al. 2019; Cai et al. 2021), we generated a phylogenomic dataset of 2,135 loci (referred to as G2135 hereafter; Data S1), each containing at least 10 out of 45 species in Malpighiales (Tables S1-2). A phylogenetic based approach from Yang and Smith (2014) was applied to remove paralogs and potential HGTs (Fig. S1). These 45 species included all three genera of Rafflesiaceae (Sapria, Rafflesia, and Rhizanthes), both genera from Apodanthaceae (Apodanthes and Pilostyles), and three outgroup species from Oxalidales and Celastrales. Individual DNA and protein alignments were filtered by the profile hidden Markov models implemented in HmmCleaner (Di Franco et al. 2019). The final concatenated matrix consisted of 3,099,966 DNA codon sites (1,033,322 protein sites) with a gene and character occupancy of 70.3% and 75.5%, respectively. Each locus on average contained 32 species and the median length of DNA alignment was 1,260 bp after trimming. The mean branch support for maximum likelihood (ML) gene trees was 72.0 based on ultrafast bootstrap replicates (UFBP) (Hoang et al. 2018), but the mean support was only 57.4 UFBP for the placement of Rafflesiaceae and 36.7 UFBP for Apodanthaceae (Table S2). In addition to this complete dataset, we compiled two subsampled datasets with 446 (G446) and 829 loci (G829). The G446 dataset contained loci where Rafflesiaceae and Apodanthaceae were monophyletic in individual gene trees (Table S3; Data S1). The G829 dataset was a subset of lock-like loci suitable for divergence time estimation and the application of computationally intensive mixture models (Data S1). See Methods below for details on subsampling criteria. All curated alignments and gene trees are available as supplementary data on Zenodo (10.5281/zenodo.11643025).

### Identifying suitable genes for placing Rafflesiaceae

To identify suitable genes for phylogenetic reconstruction, especially regarding the placement of Rafflesiaceae and Apodanthaceae, we used fourteen metrics to quantify various aspects of phylogenetic properties (Table S2). These metrics included six tree-wise features such as number of species, number of parsimony informative sites, average branch support, total tree length, root-to-tip distance variance, and gene tree–species tree congruence; and eight metrics specific to Rafflesiaceae and Apodanthaceae, including their DNA compositional bias, lineage-specific branch lengths, and branch support. These fourteen metrics were significantly correlated in 50 out of the 91 comparisons (Pearson correlation p-value <0.05; Fig. S2). For example, the mean branch support, which is a widely applied metric for selecting high-quality phylogenetic markers, was positively correlated with the number of parsimony informative sites (p-value = 8.9e-16; Pearson correlation coefficient = 0.225) and gene tree–species tree congruence (p-value = 0; Pearson correlation coefficient = 0.303). Rafflesiaceae- and Apodanthaceae-specific features were strongly correlated with each other, but relatively independent of the tree-wise metrics. For example, the branch support of Rafflesiaceae and Apodanthaceae was only significantly positively correlated with each other and their branch lengths, but not any tree-wise features (Fig. S2). The only exceptions involved the correlated branch length between Rafflesiaceae, Apodanthaceae, and the total tree length and variation because the long terminal branches of these parasites had a substantial influence on the global mean.

Initial phylogenetic reconstructions yielded weakly supported species trees Explorative phylogenetic reconstructions using concatenation and coalescent methods unanimously supported the monophyly of Rafflesiaceae and Apodanthaceae (100 UFBP or 1.0 posterior probability (PP) in all 22 analyses). Despite the effort to mitigate homoplasy, rate heterogeneity, and ILS, the relative replacement of Rafflesiaceae and Apodanthaceae within Malpighiales remained elusive (Table 1; Table S4; Data S2).

**Table 1.** Phylogenetic analysis schemes for 22 concatenation and coalescent methods. The placement and support for Rafflesiaceae and Apodanthaceae are listed. See Fig. 1B for the definition of topologies H1–H8 and branch support is evaluated by 1000 ultrafast bootstrap (UFBP) in IQ-TREE, local posterior probability (PP) in ASTRAL, and non-parametric bootstrapping in MP-EST (BP). Details of the parameter settings of each analysis, including gene and site filtering strategies, are provided in Table S4.

|  | ID | Data type | Gene set | Starting partition | Substitution model | Program | Placement | Support |
| --- | --- | --- | --- | --- | --- | --- | --- | --- |
| Concatenation | 1 | DNA | G2135 | Gene | ModelFinder | IQ-TREE | H1 | 84 UFBP |
|  | 2 | DNA | G2135 | Codon | ModelFinder | IQ-TREE | H1 | 99 UFBP |
|  | 3 | DNA | G2135 | Codon | ModelFinder | IQ-TREE | H2 | 41 UFBP |
|  | 4 | DNA | G2135 | Codon | ModelFinder | IQ-TREE | H3 | 38 UFBP |
|  | 5 | DNA | G2135 | Site | GHOST | IQ-TREE | H4 | 63 UFBP |
|  | 6 | DNA | G446 | Gene | ModelFinder | IQ-TREE | sister to all other Malpighiales | 55 UFBP |
|  | 7 | Protein | G2135 | Gene | ModelFinder | IQ-TREE | H3 | 33 UFBP |
|  | 8 | Protein | G2135 | Gene | ModelFinder | IQ-TREE | Centroplacaceae | 27 UFBP |
|  | 9 | Protein | G829 | Site | PMSF | IQ-TREE | H5 | 69 UFBP |
|  | 10 | Protein | G446 | Gene | ModelFinder | IQ-TREE | sister to all other Malpighiales | 44 UFBP |
| Coalescent | 11 | DNA | G2135 | Codon | GTR (no +I/R/G) | ASTRAL IV | H4 | 0.99 PP |
|  | 12 | DNA | G2135 | Codon | ModelFinder | ASTRAL IV | H6 | 0.48 PP |
|  | 13 | DNA | G2135 | Gene | GTR (with +I/R/G) | ASTRAL IV | H7 | 0.67 PP |
|  | 14 | DNA | G2135 | Codon | GTR (with +I/R/G) | ASTRAL IV | H8 | 0.35 PP |
|  | 15 | DNA | G2135 | Gene | ModelFinder | ASTRAL IV | H5 | 0.51 PP |
|  | 16 | DNA | G446 | Gene | ModelFinder | ASTRAL IV | H5 | 0.52 PP |
|  | 17 | DNA | G2135 | Codon | GTR (no +I/R/G) | MP-EST | H4 | 4 BP |
|  | 18 | DNA | G2135 | Codon | ModelFinder | MP-EST | H7 | 20 BP |
|  | 19 | DNA | G2135 | Gene | GTR (with +I/R/G) | MP-EST | H5 | 14 BP |
|  | 20 | DNA | G2135 | Codon | GTR (with +I/R/G) | MP-EST | H8 | 17 BP |
|  | 21 | DNA | G2135 | Gene | ModelFinder | MP-EST | H7 | 8 BP |
|  | 22 | DNA | G446 | Gene | ModelFinder | MP-EST | H5 | 7 BP |

Concatenated DNA analyses of the G2135 dataset placed Rafflesiaceae and Apodanthaceae as sister to the MRCA of Rhizophoraceae+Erythroxylaceae and Ixonanthaceae with moderate to high support (89–99 UFBP; Analysis 1 and 2 in Table 1). However, reduced support and alternative topologies were obtained when attempting to mitigate saturation by removing the third codon position or using protein sequences. For example, after removing the third codon position and applying the more stringent alignment site homology filtering criteria, Rafflesiaceae and Apodanthaceae were placed as sister to Euphorbiaceae+Peraceae with low support (33-38 UFBP) based on DNA and protein alignments (Analysis 4 and 7 in Table 1). The application of complex heterotachy and profile mixture models did not improve the result either. The DNA analysis using the GHOST heterotachy model with four mixture classes placed Rafflesiaceae and Apodanthaceae with Linaceae with a moderate support of 63 UFBP. Application of the protein profile mixture model placed Rafflesiaceae and Apodanthaceae with the MRCA of Euphorbiaceae+Peraceae, Putranjivaceae, and Pandaceae (69 UFBP). Finally, we explored the performance of an a posteriori set of 446 loci (G446) where Rafflesiaceae and Apodanthaceae were monophyletic in gene trees. Both DNA and protein analyses of G446 placed Rafflesiaceae and Apodanthaceae as sister to all other members of the Malpighiales with low support (44–55 UFBP; Analysis 6 and 10 in Table 1).

Compared to concatenation, coalescent analyses recovered more consistent results. Ten out of the twelve coalescent analyses placed Rafflesiaceae and Apodanthaceae variously with Euphorbiaceae+Peraceae, Putranjivaceae, Pandaceae, and Rhizophoraceae+ Erythroxylaceae with moderate to low support (0.35–0.99 PP; 7–20 BP; Analysis 11-22 in Table 1). Even when using the G446 dataset, both ASTRAL and MP-EST placed Rafflesiaceae and Apodanthaceae as sister to the MRCA of Putranjivaceae, Pandaceae, and Euphorbiaceae+Peraceae. However, in two analyses using the gene trees inferred from the GTR model without rate heterogeneity across sites, both ASTRAL and MP-EST placed Rafflesiaceae and Apodanthaceae with Linaceae (0.99 PP in ASTRAL and 4 BP in MP-EST). In particular, gene trees from this enforced GTR model recovered an exceptionally high proportion of monophyletic Rafflesiaceae and Apodanthaceae (n = 653) among individual gene trees compared to models allowing for rate heterogeneity across sites (n = 423–446; Table S3).

Despite these conflicts, initial analyses using concatenation and coalescent frequently placed Rafflesiaceae and Apodanthaceae with four clades consisting of Euphorbiaceae+Peraceae, Putranjivaceae, Pandaceae, and Erythroxylaceae+Rhizophoraceae. The divergence between these clades was characterized by a series of short internal branches, leading to extensive phylogenetic conflicts (Fig.1).

**Fig 1.**
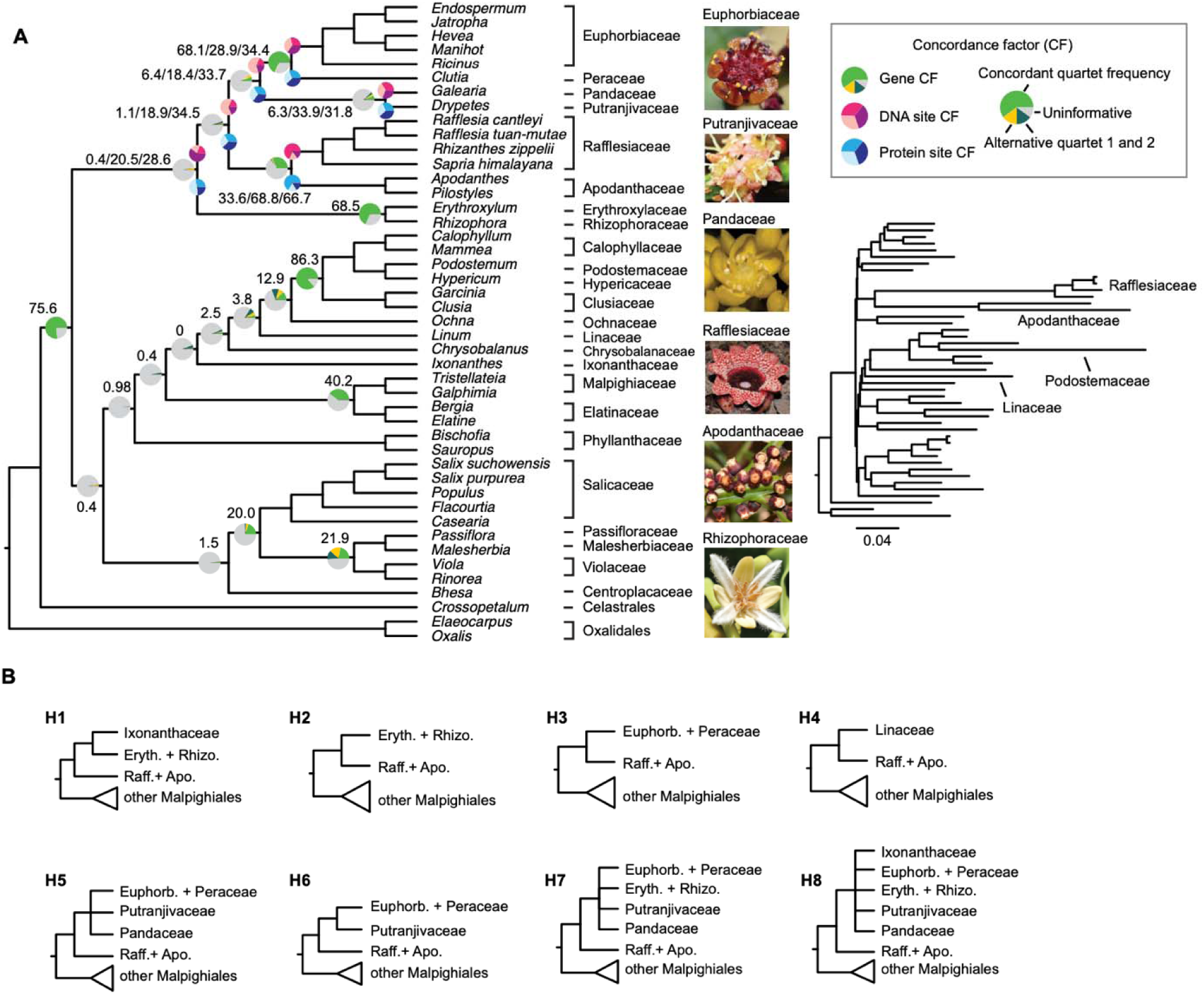
Conflicting phylogenetic signals evidenced by gene and site concordance factors lead to weakly supported conflicting placements of Rafflesiaceae and Apodanthaceae. A. The species tree cladogram is inferred by ASTRAL from the complete dataset (Analysis 15 in Table 1). The phylogram with branch lengths in mutational units shown on the right is estimated from IQ-TREE using concatenated DNA alignments. For close relatives of Rafflesiaceae and Apodanthaceae, the gene, DNA site, and protein site concordance factors (CFs) are labeled as numbers and pie charts on each node. For the rest of the tree, only gene concordance factors are labeled and visualized. B. Alternative placement of Rafflesiaceae and Apodanthaceae (Raff. + Apo.) across various data subsampling and species tree inference schemes. The 8 hypothesized placements of Rafflesiaceae and Apodanthaceae are variously supported by coalescent or concatenation analyses detailed in Table 1 and Table S4. Family names abbreviated for illustrative purposes include Rhizophoraceae (Rhizo.), Erythroxylaceae (Eryth.), and Euphorbiaceae (Euphorb.).

The concordant gene frequencies (gCF) among these four clades plus Rafflesiaceae and Apodanthaceae, defined as the percentage of decisive gene trees supporting that branch (Minh, Hahn, et al. 2020), ranged from 0.41% to 6.39%, with most genes being uninformative (86.22–96.09%). The gCFs for other early-diverging lineages in Malpighiales were similarly low and ranged from 0.02% to 3.85%, reflecting the well-recognized recalcitrant relationship in Malpighiales. In sharp contrast, internal branches within other well-recognized clades such as the parietal clade (MRCA of Salicaceae and Violaceae) and the clusioids (MRCA of Clusiaceae and Calophyllaceae) had gCFs between 20.00–86.32%.

Site concordance factors (sCF) showed similar patterns. The sCF ranged from 18.4% to 33.9% for DNA and from 28.6% to 34.5% for protein alignments among clades closely related to Rafflesiaceae and Apodanthaceae (Fig. 1). The GC content within the coding regions of Rafflesiaceae and Apodanthaceae was comparable to other free-living Malpighiales and was unlikely to be responsible for the low concordance factors observed above (Fig. S3).

### Fast-evolving genes and sites drive biased support

We evaluated the goodness-of-fit of individual genes and sites to eight alternative placements of Rafflesiaceae and Apodanthaceae derived from explorative phylogenetic analyses (defined in Table 1) using the method from Shen et al. (2017). In particular, we examined how spurious relationships might arise from fast-evolving genes and sites. The results turned out to be sensitive to codon position, substitution rate, and type of input data (i.e., DNA versus protein).

We first examined the log-likelihoods (LL) of individual sites and confirmed that the differences of log-likelihoods (ΔLL) among topologies were similar and there was no site with outstanding ΔLL values (Fig. S4). However, the site rate among the three codon positions varied substantially, with the rates in the third codon being nearly five times (median = 1.954) compared to those in the first (median = 0.483) and second codon positions (median = 0.327; Fig. S5). When evaluating topologies based on concatenated codon positions, both the first and third codons strongly rejected H5, where Rafflesiaceae and Apodanthaceae were placed with Euphorbiaceae+Peraceae, Putranjivaceae, and Pandaceae (approximately unbiased (AU) test p-value < 9.02e-56; Table S5). In contrast, the second codon supported H5 as a feasible topology (AU test p-value = 0.116; Table S5). The first codons also showed a slight preference for H1 compared to H3 (bp-RELL_H1_ = 0.641; bp-RELL_H3_ = 0.162; Table S5) whereas the third codon showed a slight preference for H3 instead (bp-RELL_H1_ = 0.187; bp-RELL_H3_ = 0.720) based on the resampling estimated log-likelihoods (RELL) (Kishino et al. 1990).

Next, we used ΔLL to identify any outlier gene showing exceptionally high support for one topology over another. For DNA alignments, the ΔLL between the best-supported topology H1 and most other alternative topologies was within a similar range (Fig. S6). However, one gene (locus ID = 313) had exceptionally large ΔLL for topology H3, H7, and H8 compared to H1 (Fig. S6), the former of which generally placed the parasites with Euphorbiaceae+Peraceae (H3) or their close relatives (H7, H8) instead of Rhizophoraceae+Erythroxylaceae and Ixonanthaceae (H1). In addition, the same gene showed a reverse trend by strongly favoring H3 over H1 based on protein alignment. We thus removed this gene before investigating the impact of substitution rate on topology support.

To further explore how fast-evolving genes prone to long-branch attraction may contribute to the conflicting placements of Rafflesiaceae and Apodanthaceae, we ordered genes based on the substitution rate estimated by IQ-TREE, which ranged from 0.523 to 1.897 for DNA alignments and 0.078 to 3.056 for protein alignments. We examined how the support for optimal topology changed with the addition of genes with increasing rates. As a negative control where a topology is universally rejected regardless of evolutionary rates, we compared the ΔLL of the optimal topology (H1 in DNA alignments and H3 in protein alignments) to H4, which placed the parasites with Linaceae. Both DNA and protein data demonstrated a consistent and nearly linear increase in ΔLL as more rapidly evolving genes were added (Fig. 2C, F). The trend also aligned with the simulated mean where genes were randomly ordered regardless of rates. The decisive lead in the final ΔLL (>1000) plus the alignment between the empirical and simulated curve of ΔLL demonstrated an example when the alternative topology (H4) could be rejected with confidence (AU test p-value <0.002; Table S5) and the impact of substitution rate was minimal. In contrast, support for H1 (Rhizophoraceae+Erythroxylaceae, Ixonanthaceae) over H3 (Euphorbiaceae+Peraceae) in DNA alignments was clearly driven by a small group of fast-evolving genes (Fig. 2A, B).

**Fig. 2.**
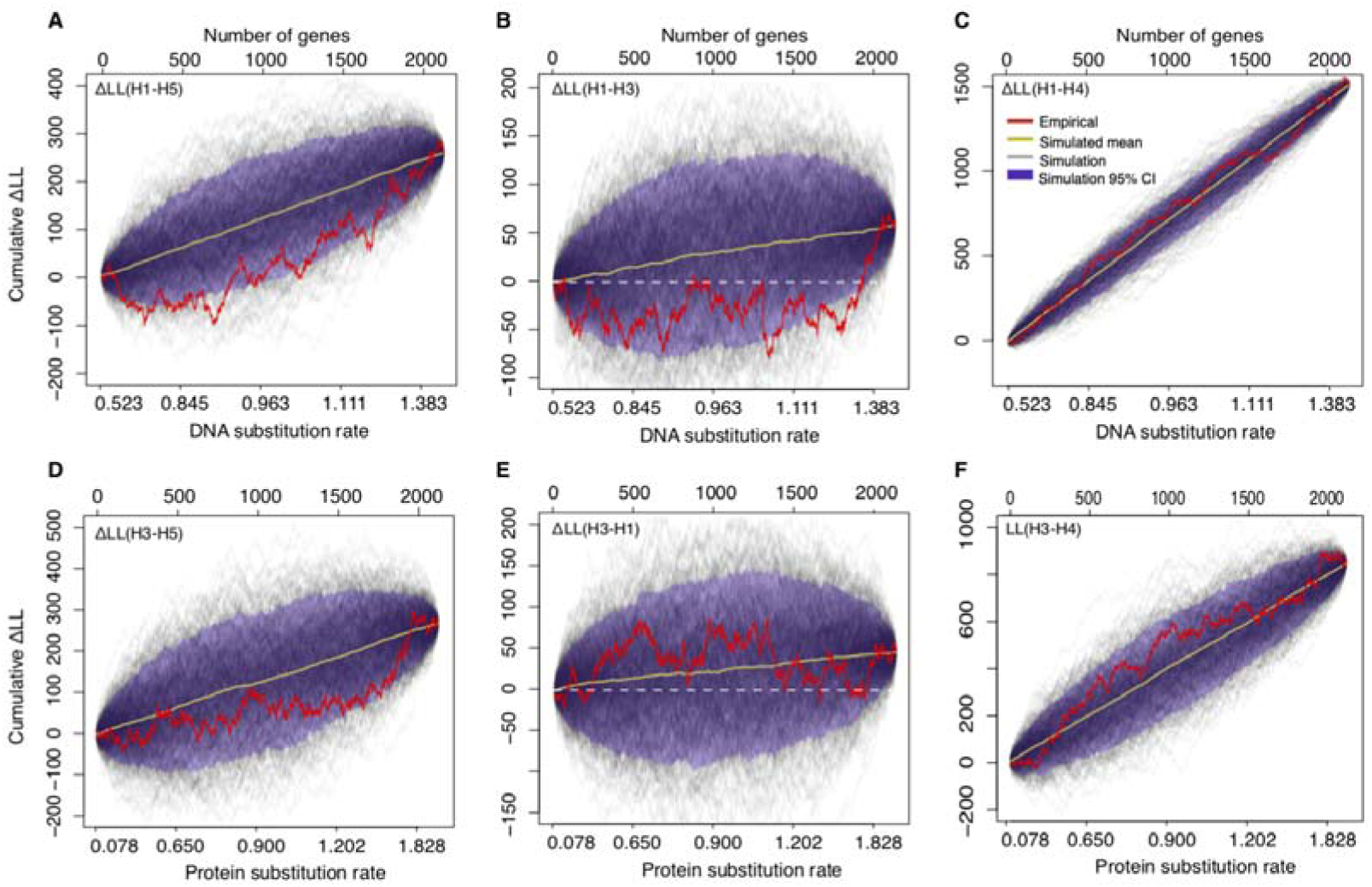
Changes in cumulative differences of log-likelihood (ΔLL) when adding more rapidly evolving genes. A–C ΔLL of the optimum topology H1 compared to three alternative topologies H5 (A), H3 (B), and H4 (C) based on DNA sequences. D–F ΔLL of the optimum topology H3 compared to three alternative topologies H5 (D), H1 (E), and H4 (F) based on protein sequences. Topologies H1-H5 are defined in Fig. 1B. All 2,135 genes are ordered on the x-axis based on DNA (A-C) or protein substitution rates (D-E). The rate and likelihoods for individual genes were inferred using the –wpr and -wpl flags under the same optimal substitution model determined by ModelFinder and per-gene partition in IQ-TREE. The red line represents the cumulative ΔLL as genes with higher rates are added. The dark gray lines represent the cumulative ΔLL from 1000 simulated bootstrap replications where genes are randomly ordered. The yellow line represents the simulated average. The gene and site rate and likelihood estimates supporting this figure are provided in Data S3.

The cumulative ΔLL of H1–H3 was below zero or near zero for the first 1,735 (81.2%) slower-evolving genes based on DNA alignments (Fig. 2B). However, the 9.8% fastest- evolving genes significantly favored the H1 hypothesis over H3 (AU test p-value = 0.147 versus 0.0114; Table S8) such that they completely reverse the trend (Fig. 2B). This phenomenon was mitigated in protein-based analyses where the empirical curve of ΔLL aligned better with the simulated average (Fig. 2E). But for both DNA and protein data, slower-evolving genes seem to support a closer affinity between the parasites and Euphorbiaceae+Peraceae (H3) instead of Rhizophoraceae+Erythroxylaceae, Ixonanthaceae (H1). Finally, we examined the changes of ΔLL regarding the H5 topology (Euphorbiaceae+Peraceae, Putranjivaceae, Pandaceae), which is favored by many coalescent analyses and one protein-based concatenation analysis using the profile mixture model. DNA alignments from slower-evolving genes similarly showed a preference for this alternative topology H5 than H1 compared to fast-evolving genes (Fig. 2A). Although this is less obvious in the protein analysis (Fig. 2D).

### The anomaly zone and coalescent simulations

To explore the effects of ILS, we first calculated the branch length in coalescent units for focal clades. The three successive short internal branches among the parasites and the other four focal clades ranged from 0.015 to 0.096 in coalescent units (Fig. 3A). The estimated minimum branch length to escape the anomaly zone was 0.344, which is four times longer than the actual branch lengths (Fig. 3B). This result suggests that these rapid radiations involving Rafflesiaceae+Apodanthaceae are clearly subject to the anomaly zone. Our gene tree simulation with six taxa also corroborated this conclusion; the most common gene tree was discordant from the true species tree (Fig. 3C), which ranked 11^th^ in frequency. The highly consistent frequencies between the empirical and simulated gene trees suggested that most gene tree variations can be explained by ILS. However, a significant difference in gene tree frequencies was observed in topology T15, in which Rafflesiaceae was placed sister to all other Malpighiales (Fig. S7). This topology was present in only 11% of the true gene trees in the simulation but increased to 17% when adding in the ML inference step (Fig. 3C).

**Fig 3.**
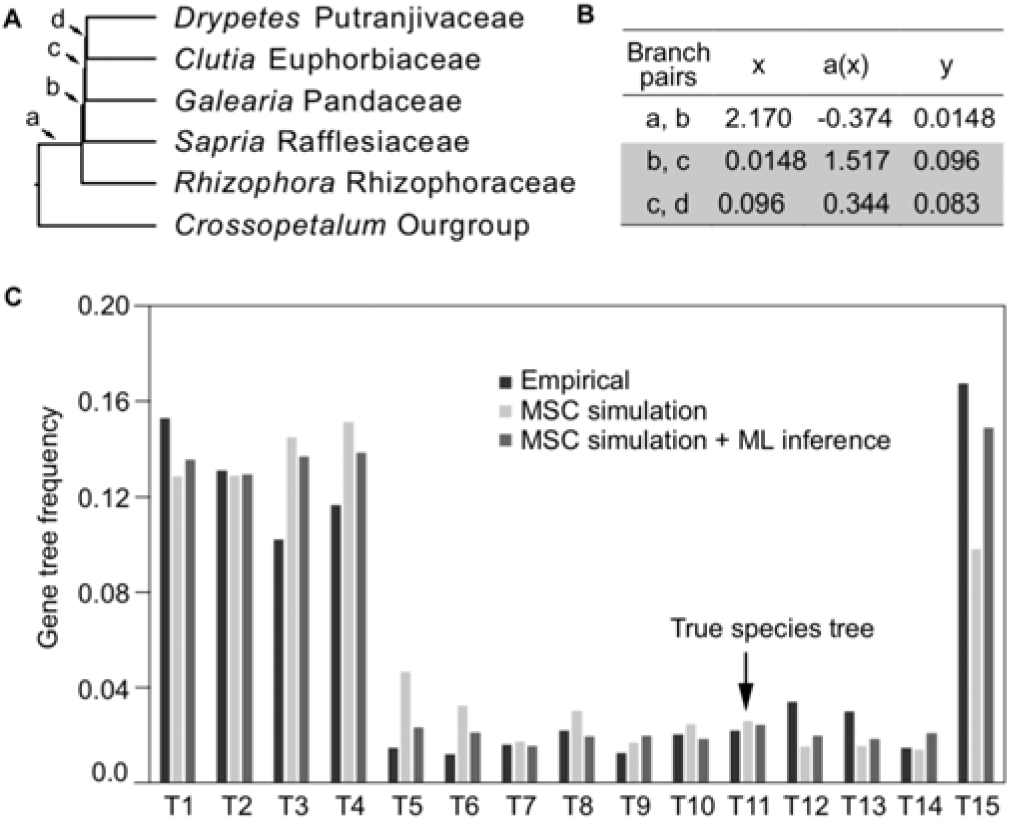
The anomaly zone and coalescent simulation. A. Reference species tree for coalescent simulation with labeled branches. B. For each internal branch pair in a four-taxon tree, boundaries of the anomaly zone a(x) calculated from (Degnan and Rosenberg 2006) are compared to the empirical branch length (y). Two out of three branch pairs (b-c and c-d) fall within the anomaly zone as y< a(x). C. Gene tree frequency for the 15 alternative placements of Rafflesiaceae (T1–T15) in the empirical gene trees (dark grey), simulated true gene trees under the multispecies coalescent model (MSC; medium grey), and inferred maximum likelihood (ML) gene trees based on the simulated sequences (light grey).

Topologies for T1 to T15 are defined in Fig. S7. In all three cases, gene trees consistent with the true species tree topology have very low frequency.

When inferring the species tree from simulated alignments, none of the concatenated analyses recovered the true species tree. Specifically, when more than 4,000 loci were used, the true topology was strongly rejected by the AU test, suggesting statistical inconsistency (Table S6). Instead, the incorrect species topologies were variously supported by AU tests. T5 and T15 were most frequently favored, which placed Rafflesiaceae with (Putranjivaceae, Euphorbiaceae) or the MRCA of Rhizophoraceae and Euphorbiaceae, respectively. In contrast, summary coalescent methods consistently recovered the true species when at least 4,000 loci were used (Table S7). The gene tree estimation error was as high as 78.1% for 500-bp-alignments and 70.0% for 2000-bp-alignments in terms of the placement of Rafflesiaceae. Such high estimation error led to incorrect species tree estimation when less than 3,000 genes were used. But even when enough loci were used and the true species tree was recovered, the BP support for Rafflesiaceae was still moderate to low (35–84 BP) and did not increase with the number of genes used. Moreover, alternative topologies could not be rejected based on the likelihood ratio test. For example, the difference in pseudolikelihoods between the true species tree H11 and an alternative topology H5 was 0.71 when using 1,000 genes (pseudolikelihood = -10968.5) and 20.3 when using 10,000 genes (pseudolikelihood = -109763.3), which were not statistically significant (p– value >0.51, Table S7).

### Divergence time estimation

Finally, based on a species tree topology from the ASTRAL analysis, we inferred the divergence time of Malpighiales using the Bayesian method implemented in PhyloBayes (Lartillot et al. 2009) and the penalized likelihood method implemented in treePL (Smith and O’Meara 2012). Both approaches supported a mid-Cretaceous origin of Malpighiales, followed by radiation in the Albian that gave rise to the majority of the family-level diversity within Malpighiales (Fig. 4; Fig. S8). The Phylobayes analysis was based on a subset of 829 clock-like loci with the third codon position removed. Crown group Malpighiales was inferred to originate during the Albian at 111.1 Ma (95% confidence interval CI = 109.5–112.1; Fig. 4) and the MRCA of Rafflesiaceae and Apodanthaceae diverged from its sister clade, the MRCA of Euphorbiaceae and Pandaceae, during the Albian at 110.9 Ma (95% CI = 109.2–111.9). Rafflesiaceae diverged from Apodanthaceae at 106.5 Ma (95% CI = 102.8–108.7) and crown group Rafflesiaceae arose very recently during the Miocene at 15.1 Ma (95% CI = 9.2–28.1). Crown group Apodanthaceae was inferred to originate in the early Eocene at 53.1 Ma (95% CI = 35.9–66.4). The treePL analysis was based on the concatenated DNA alignment of the complete G2135 dataset without the third codon. The MRCA of Rafflesiaceae and Apodanthaceae diverged from its sister clade at 106.4 Ma (Fig. S8). Stem group Rafflesiaceae diverged from Apodanthaceae in the Cenomanian at 98.8 Ma and its crown group age is inferred to be 19.0 Ma. Crown group Apodanthaceae, on the other hand, originated around the Paleocene-Eocene transition at 55.6 Ma.

**Fig. 4.**
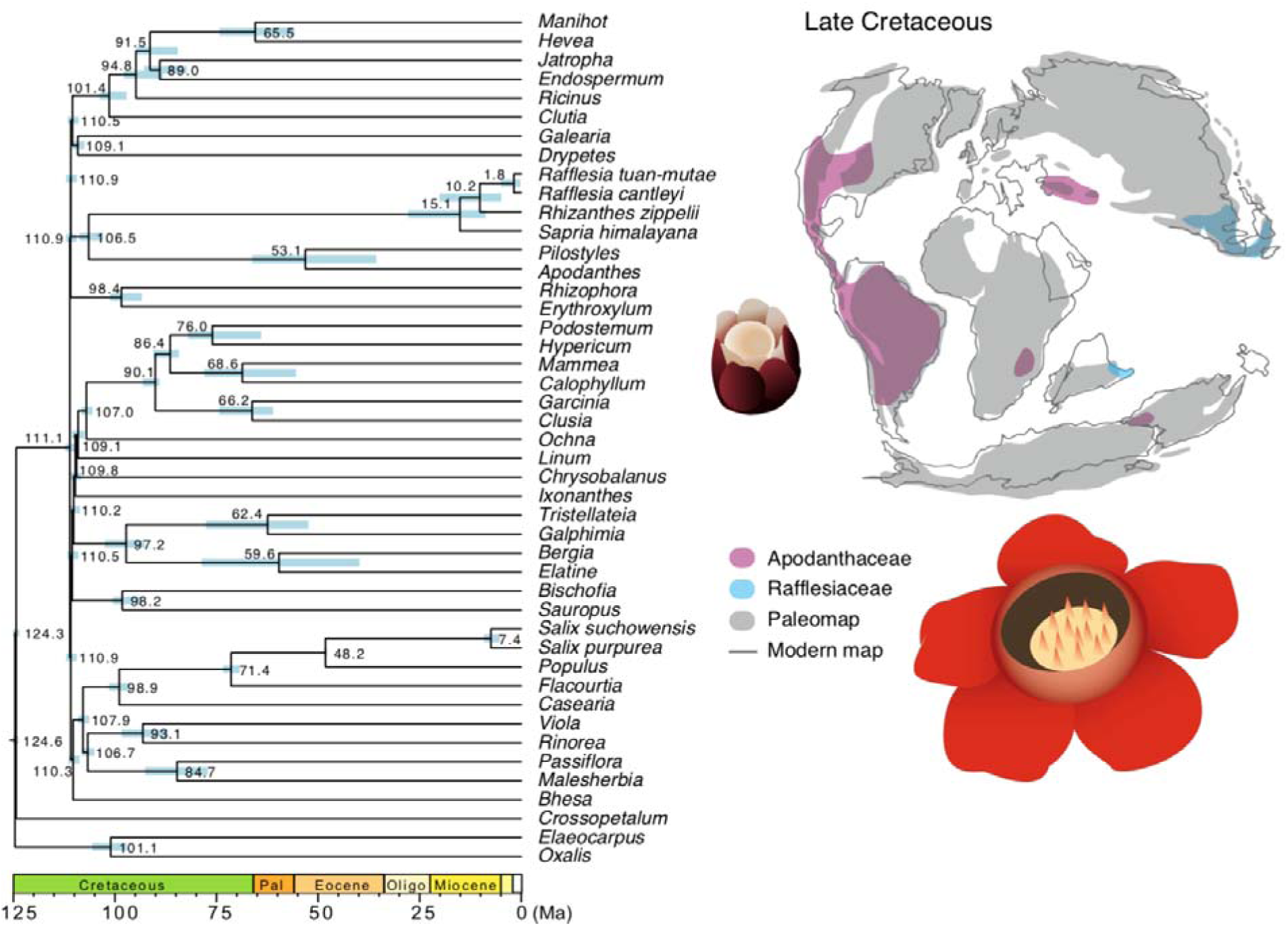
Bayesian divergence time estimation of Malpighiales. The dated phylogeny is inferred under the CAT-GTR model based on the concatenated DNA sequences from 829 clock-like genes in PhyloBayes. Divergence time measured in million years (Ma) is labeled on each node and the 95% confidence intervals are visualized as bars. Distribution of the extant species in Apodanthaceae and Rafflesiaceae is projected to the paleomap from the Late Cretaceous at 94 million years ago. The map is referenced from the PALEOMAP Project by C. R. Scotese (www.scotese.com).

## DISCUSSION

Genome-wide rate acceleration in parasites contributes to long-branch attraction Despite meticulously curated genomic data, we did not obtain a single tidy placement for Rafflesiaceae and Apodanthaceae within Malpighiales. The shift to such an extreme parasitic lifestyle early in their evolution has undoubtedly reshaped the selective forces acting on otherwise conserved autotrophic functions such as photosynthesis (Cai 2023). As a result, relaxed selection and accelerated nucleotide substitution rates are often characteristic of the plastid and nuclear genomes of these species (Bromham et al. 2013; Wicke et al. 2016; Chen et al. 2023). This rate acceleration has contributed to exceptionally long branches in Rafflesiaceae and Apodanthaceae (Fig. 1A). Previous genomic investigations in Rafflesiaceae revealed that its synonymous substitution rate was 4.6 times higher than its close relative Manihot (Euphorbiaceae) and 3.3 times higher than Populus (Salicaceae) (Cai et al. 2021). After the divergence of Rafflesiaceae from its free-living relatives, 35.1% of genes have experienced more than one synonymous substitution per site (Cai et al. 2021) and this has led to extensive phylogenetic conflicts when trying to place these parasites. These conflicts are reflected by the low gene and site concordance factors where a remarkable 98.9% of the genes are uninformative for placing Rafflesiaceae and Apodanthaceae (Fig. 1).

Compared to parsimony, likelihood methods are more robust to long-branch attractions (Felsenstein 1978; Huelsenbeck and Hillis 1993), but overly simplistic substitution models can still lead to long-branch attraction (Hendy and Penny 1989; Zharkikh and Li 1993; Tateno et al. 1994; Bruno and Halpern 1999; Ho and Jermiin 2004; Bergsten 2005; Brinkmann et al. 2005; Philippe et al. 2011). In Malpighiales, models without sufficient rate heterogeneity accommodation often grouped Rafflesiaceae and Apodanthaceae with other apparent “rogue” taxa with long branches such as Linaceae (Analysis 5, 11, and 17 in Table 1). For example, the concatenation Analysis 5 employed the GHOST heterotachy model but our high-performance computing cluster can only accommodate four mixture classes due to its high computational demand. As a result, despite designed to mitigate heterotachy, the GHOST model with four mixture classes appeared to be inadequate for capturing cross-site rate variation compared to partition models with 107 to 178 partitions, each with different rate heterogeneity modeling. This overly simplified substitution model in our GHOST analysis may have contributed to the grouping of Linaceae, Rafflesiaceae, and Apodanthaceae, all of which have long root-to-tip branch lengths. In contrast, despite the even longer terminal branch in Podostemaceae and its high GC content, the phylogenetic affinity between Podostemaceae and Hypericaceae was maximally supported in all analyses due to the longer internal branch supporting their shared ancestry (Fig. 2A; Fig. S3; Data S2).

Our strongest evidence for the influence of long branch attraction stems from the a posteriori selection of genes and models where Rafflesiaceae and Apodanthaceae are monophyletic. Concatenation of the G446 dataset where the parasites were monophyletic in gene trees placed Rafflesiaceae and Apodanthaceae sister to all other Malpighiales, which is a classic signal of long branch attraction (Kapli et al. 2020). Coalescent analyses of the same set of genes, on the other hand, recovered a more conventional topology (H5).

This result suggests that coalescent methods might be more robust to long branch attraction compared to concatenation methods, but they are still not immune to these effects. Among the five gene tree reconstruction methods, applying the GTR model with no rate heterogeneity modeling yielded up to 54.3% more monophyletic Rafflesiaceae and Apodanthaceae across genes (Table S3), but coalescent analysis of these gene trees using ASTRAL and MP-EST consistently placed the parasites with Linaceae, similar to the concatenation results above using the oversimplified GHOST model (Fig. 2C, F). Many of these outcomes of long branch attraction were captured by our simulation. For example, gene tree frequencies for the fifteen alternative topologies were largely consistent between the true gene trees and gene trees inferred from the simulated alignments (Fig. 3C). Yet a disproportionately high frequency of T15 placing Rafflesiaceae sister to all other Malpighiales was recovered among ML-inferred gene trees compared to the true gene trees, likely a result of long-branch attraction during the ML inference step. When aggregating such erroneous signals into concatenated alignments, it contributed to the support of T15 in optimum species tree searches and rejection of the true species tree (Fig. 2B; Table S6).

Navigating the phylogenetic “Danger Zone” of excessive incomplete lineage sorting and long-branch attraction

Like many other rapid radiations in mammals and birds (Liu et al. 2017; Stiller et al. 2024), the explosive malpighialean speciation followed by parasitism-induced rate acceleration created a sharp contrast in branch length before and after the origin of stem group Rafflesiaceae+Apodanthaceae. The combination of high ILS and long-branch attraction presents significant challenges for species reconstruction in both concatenation and coalescent methods (Martyn and Steel 2012; Roch et al. 2019). We define the term “danger zone” to represent an empirically identified species tree space under the extreme influence of ILS and long-branch attraction such that phylogenetic reconstruction is excessively challenging.

Selecting reliable genes with desired phylogenetic properties under these circumstances is one of the ways to optimize signal-to-noise ratio (Koch 2021) and potentially deal with the “danger zone” problem. Here, we identified that some commonly applied gene-filtering criteria, including high gene tree–species tree congruence, high gene tree support, and low missing taxa, strongly covary in our dataset (Fig. S1). Yet none of these global metrics were useful when choosing informative genes to place Rafflesiaceae and Apodanthaceae. For example, the branch support of Rafflesiaceae and Apodanthaceae was only strongly influenced by the length of its short stem group branch rather than the long descending branch leading to extant genera (Fig. S1). Therefore, the recalcitrant placement of Rafflesiaceae and Apodanthaceae appears to be primarily driven by incomplete lineage sorting. But long-branch attraction caused by fast evolving genes and sites further exacerbated the problem and led to spurious relationships (Fig. 2). Our application of the G446 dataset where Rafflesiaceae and Apodanthaceae were monophyletic was aimed to select “informative” loci based on a posteriori criterion, but counterintuitively, phylogenetic reconstruction using these genes demonstrated that they were more prone to long branch attraction (Table 1). Thus, selecting a small set of highly informative genes is not a promising approach to address the “danger zone” problem because an empirically verifiable criterion to optimize signal-to-noise ratio is lacking in our system and many others.

Alternatively, other efforts to resolve clades similarly suffering from “danger zone” phenomena have turned to a big-data analytics approach. Examples include the application of tens of thousands of loci from whole genome sequencing to resolve the bird Tree of Life (Stiller et al. 2024). Yet our simulation concluded that current methods for modeling substitution and coalescent processes are insufficient to fully resolve the placement of Rafflesiaceae and Apodanthaceae with at least ten thousand loci. The empirical species tree existed within a parameter space where both incomplete lineage sorting and long-branch attraction were prevalent. Concordant gene trees appeared at a very low frequency, and the most common gene trees consistently placed the parasite sister to all other Malpighiales due to long-branch attraction (Fig. 3C). In addition, massive gene loss demonstrated in Rafflesiaceae renders a greatly reduced pool of phylogenetic markers that are potentially insufficient to achieve a definitive placement. Alternative methods employing macro- and micro-scale gene synteny and rare genomic changes have been increasingly implemented to untangle recalcitrant relationships in angiosperms and teleost fishes (Zhao et al. 2021; Parey et al. 2023). As more genomic data is developed in Malpighiales and beyond, the application of these new methods may provide novel insights into this thorny phylogenetic problem.

The Gondwana origin of Rafflesiaceae and Apodanthaceae

Despite the phylogenetic uncertainty, coalescent analyses and concatenation analyses with complex models consistently recover similar sister clades for Rafflesiaceae and Apodanthaceae. This improvement in topological resolution, plus the mid-Cretaceous divergence time of stem group Rafflesiaceae and Apodanthaceae, has led to novel insights into the early evolutionary history of these enigmatic parasites.

Most coalescent and mixture-model-based concatenation analyses placed Rafflesiaceae and Apodanthaceae with clades variously consisting of Putranjivaceae, Pandaceae, Euphorbiaceae, and Peraceae (Table 1), all of which share a pantropical distribution.

Although the highly modified morphology of Rafflesiaceae and Apodanthaceae makes it challenging to identify synapomorphies, several shared features support such phylogenetic affinity. For example, the pentamerous dioecious flowers in Rafflesiaceae and Apodanthaceae are commonly found in Pandaceae, Putranjivaceae, and its unsampled close relative Lophopyxidaceae (Matthews and Endress 2013; Wurdack et al. 2004). In addition, Putranjivaceae releases sulfur-rich, strong-smelling chemicals to attract pollinators like Rafflesiaceae and Apodanthaceae. The foul odor of Rafflesiaceae largely comes from dimethyl disulfide and dimethyl trisulfide (Jürgens et al. 2013; Wee et al. 2018), whereas the strong smell in Drypetes (Putranjivaceae) comes from isothiocyanates of the glucosinolates pathway (Johnson et al. 2009). Future studies focusing on the metabolic pathway of these chemicals may elucidate whether they are derived from shared ancestry.

The parasitic common ancestor of Rafflesiaceae and Apodanthaceae diverged from their free-living relatives in the mid-Cretaceous as part of the initial malpighialean radiation (110.9 Ma; 95% CI = 109.2–111.9). This was then followed by the divergence between stem group Rafflesiaceae and Apodanthaceae at 106.5 Ma (95% CI = 102.8–108.7). These Albian age estimates not only suggested that extreme life history strategies such as endoparasitism can evolve rapidly within less than 5 Ma but also pinpointed Rafflesiaceae and Apodanthaceae as the oldest endoparasitic and holoparasitic clades known. Many other holoparasitic lineages such as Hydnoraceae and Cynomoriaceae have comparable stem group ages in the mid-Cretaceous (Naumann et al. 2013), but their much younger or unknown crown group ages make it difficult to determine the exact date when holoparasitism evolved. Our estimation of the crown group Apodanthaceae in Eocene (53.1 Ma, 95% CI = 35.9–66.4) aligns with those from Naumann et al. (2013) and Bellot and Renner (2014), but we drastically push forward the age of crown group Rafflesiaceae to Miocene (15.1 Ma; 95% CI = 9.2–28.1). This Miocene age is the youngest compared to previous estimates from Bendiksby et al. (2010) at 81.7 Ma, Pelser et al. (2019) at 68 Ma, and Naumann et al. (2013) at 65.3 Ma; but it aligns well with the relative age constraint imposed by horizontal gene transfer (Davín et al. 2018)—the crown group age of the recipient (Rafflesiaceae) should be younger than the stem group age of the donor (Ampelopsis and Tetrastigma; 41.2-50.6 Ma) (Chen et al. 2011; Nie et al. 2012a; Cai et al. 2021). We anticipate that these age estimates will continue to be refined in future studies as more comprehensive taxon sampling, fossil calibrations, and branch-rate modeling are applied, such as those implemented in starBEAST (Ogilvie et al. 2017).

The mid-Cretaceous divergence and the distribution of extant Rafflesiaceae and Apodanthaceae in South America, Africa, Australia, and Northern India strongly suggest a Gondwanan origin (Fig. 4). Such a fragmented global distribution makes likelihood-based biogeographic reconstruction untraceable. Given the poor long-distance dispersal capacity and restricted current range of these endoparasites (Barkman et al. 2017; Malabrigo Jr. et al. 2025), we propose a southern Gondwana origin and an “out-of-India” dispersal route for Rafflesiaceae. Extant Rafflesiaceae are frequently found in southern Thailand to the Philippines, but the early-diverging Sapria himalayana are distributed in the seasonal forest south of the Himalayas in Assam India and Myanmar. This potentially points to an Indian origin of stem group Rafflesiaceae that rafted the continent until its collision with Eurasia during the Eocene. Particularly, we previously demonstrated a host shift in stem group Rafflesiaceae from Ampelopsis to Tetrastigma (Cai et al. 2021). The major geographical and climatic transitions associated with the India collision may well have served as a trigger for such shifts in life history strategies. The oldest fruit fossil of Vitaceae from the Late Cretaceous was also discovered in India (Manchester et al. 2013). Similar “out-of-India” dispersal has been demonstrated in Dipterocarpus, Crypteroniaceae, and multiple lineages of frogs (Bossuyt and Milinkovitch 2001; Conti et al. 2002; Bansal et al. 2022). A northern route via Laurasia is also possible during the Paleocene and Eocene, as has been proposed for the historical host clade Ampelopsis (Nie et al. 2012b). Future fossil discoveries, especially involving the distinctive seeds of Rafflesiaceae and Apodanthaceae, may help resolve this mystery.

On the other hand, the late Paleocene to early Eocene divergence between Apodanthes and Pilostyles in crown group Apodanthaceae was likely caused by ecological- and host-related shifts. This divergence time marks the warmest epoch on earth that permitted the dispersal of dry tropical Pilostyles through southern Patagonia, Antarctica to Australia, the corridor of which did not close until late Eocene (Amoo et al. 2022). However, our Eocene age estimates necessitate repeated long-distance dispersal of Pilostyles from the neotropics to tropical Africa and Persia. The patchy distribution of Pilostyles in present-day Persia that corresponds to the ancient Cimmerian continent could have been caused by large-scale extinction events due to major volcanism events and drastic paleoclimate shifts during the repositioning of this plate.

## Conclusion

Our empirical investigation and simulation suggested that the combination of excessive incomplete lineage sorting and long-branch attraction created a seemingly intractable phylogenetic “danger zone” where the placement of Rafflesiaceae and Apodanthaceae may never be resolved neatly. The unique life-history strategy of these endoparasites has led to high substitution rates that contribute to long-branch attraction in both concatenation and coalescent analyses. Potential sister clades of Rafflesiaceae and Apodanthaceae include Putranjivaceae, Pandaceae, Euphorbiaceae, and Peraceae, all sharing a pantropical distribution. Divergence time estimates suggest that Rafflesiaceae and Apodanthaceae represent the oldest known parasitic plant lineage, with plant endoparasitism evolving in the ecosystem as early as the mid-Cretaceous. The divergence between Rafflesiaceae and Apodanthaceae and the early diversification of Pilostyles (Apodanthaceae) within the neotropics and southern Australia may have been influenced by the Gondwana vicariance. These novel phylogenetic and temporal insights have opened a series of questions regarding the processes that shaped the distinct floral size, geographical distribution, and host preference in these two families. In particular, what may serve as the ancestral host(s) for Rafflesiaceae and Apodanthaceae given their highly specific present-day host preference in Vitaceae, Salicaceae, and Fabaceae? Perhaps close examination of host-derived gene transfers will help reveal the hidden history of these remarkable plants in deep time.

## Materials and Methods

### RNA sequencing and transcriptome assembly of Apodanthaceae

Apodanthaceae was recently demonstrated to be a close relative of Rafflesiaceae (Alzate et al. 2024) and we thus obtained the transcriptomes of both Apodanthaceae genera for phylogenomic investigation. Sample collection, RNA extraction, sequencing, and transcriptome assembly of Pilostyles boyacensis and Apodanthes caseariae were described in González et al. (2020) and Alzate et al. (2024), respectively. Briefly, for each species, three tissue samples spanning the full life cycle, including endophytic tissue, preanthetic flowers, and fruits and seeds, were flash frozen in the field. The Pilostyles samples were at xerophytic thickets around Villa de Leyva (Boyacá, Colombia) and voucher specimens were deposited at COL (voucher FG 4518 and voucher FG 4519). The Apodanthes samples were collected in wet tropical forests around Villavicencio (Department of Meta, Colombia) and the voucher specimen was deposited at HUA (voucher NP 495).

For RNA extraction, we applied the TRIzol_TM Reagent (Invitrogen) Kit and the Spectrum Plant Total RNA Kit (Sigma-Aldrich TM) for Pilostyles and Apodanthes, respectively. RNA sequencing library was prepared using the TruSeq Stranded mRNA library construction kit (Illumina) and sequenced on the Illumina Novaseq 6000 instrument in Macrogen (South Korea). Read cleaning was performed with a quality threshold of Q30 and a minimum read length of 70 bp, singletons were excluded. Contig was assembled using the Trinity V2.5.1 software (Grabherr et al. 2011) with TRIMMOMATIC adapter removal.

### Phylogenomic dataset assembly

We combined the phylogenomic datasets from our previous studies (Cai et al. 2019; Cai et al. 2021) and transcriptomes from Apodanthaceae to generate a 2,135-gene dataset (Data S1). The combined dataset included 45 species representing all five extant genera of Rafflesiaceae and Apodanthaceae, twenty-one families broadly sampled across Malpighiales, and three outgroup species from Oxalidales and Celastrales (Table S1).

Briefly, the two datasets from Cai et al. (2019) and Cai et al. (2021) were merged based on gene IDs of the three Malpighiales species sampled in both studies (Manihot esculenta, Populus trichocarpa, and Jatropha curcas). Preliminary DNA alignments and gene trees were inferred for each orthogroup using MAFFT v.7.299 (Katoh and Standley 2013) and FastTree v2.1 (Price et al. 2010). These preliminarily filtered alignments were used to build profile hidden Markov models to search for orthologous sequences in Apodanthaceae using HMMER v3.4 (Mistry et al. 2013). For each ortholog, only the best HMMER hit with e-values lower than 1e-70 was used for Apodanthes and Pilostyles. Finally, the tree-based ortholog identification approach from Yang and Smith (2014) was applied iteratively to remove HGT genes in Rafflesiaceae and Apodanthaceae as well as paralogs arising from gene duplications. Any taxa with branches longer than 1.5 were removed. This cutoff was determined by the global distribution of branch length where 1.5 is a natural break (Fig. S9). The final phylogenomic dataset consisted of 2,135 protein-coding orthogroups satisfying the following three criteria: (1) included at least one vertically transmitted gene from Rafflesiaceae or Apodanthaceae; (2) included at least 10 species (23% species completeness); (3) all genes are orthologous to each other, and no paralogs generated by gene duplications prior to the divergence of Malpighiales were included. A detailed description of dataset assembly is provided in the supplementary note S1. All scripts used in this study are available on the GitHub repository (https://github.com/lmcai/Rafflesiaceae_phylogenomics).

### Sequence alignment, cleaning, and gene tree reconstruction

Protein sequences of these 2,135 single-copy orthogroups were aligned using the E-INS-i algorithm implemented in MAFFT and then converted into the corresponding codon alignments using pal2nal (Suyama et al. 2006). Protein and DNA alignments were filtered using the profile hidden Markov models implemented in HmmCleaner v0.180750 (Di Franco et al. 2019). We applied a threshold of 50 and 10 to generate two sets of alignments with relaxed or stringent site filtering, respectively. In particular, a threshold of 50 can effectively remove nonhomologous regions while retaining fast-evolving sequences in Rafflesiaceae and Apodanthaceae. Consequently, 0.67% of the nucleotides and 0.94% of the amino acids from the raw alignments were flagged. In contrast, a threshold of 10 will remove these fast-evolving sequences and 2.4% of the nucleotides and 4.4% of the amino acids were flagged. We then used a custom Python script to mask the flagged regions and remove sites with >70% ambiguous characters (i.e., gaps and masked sites) while retaining codon positions for downstream analysis (available on the GitHub repository).

We inferred ML phylogenies for each locus based on five combinations of DNA partitioning strategies and substitution models using IQ-TREE v2.3.6 (Minh, Schmidt, et al. 2020). These ML inferences included (1) codon-based partition and forcing the GTR substitution model without rate heterogeneity across sites (i.e., no +I or +R or +G); (2) codon-based partition and forcing the GTR substitution model with rate heterogeneity across sites; (3) codon-based partition and using the optimal substitution models determined by ModelFinder (Kalyaanamoorthy et al. 2017); (4) no partition within loci and forcing the GTR substitution model with rate heterogeneity across sites; (5) no partition within loci and using the optimal substitution models determined by ModelFinder. For each ML analysis, branch supports were assessed using ultrafast bootstrap approximation (UFBP) (Hoang et al. 2018) with 1000 replicates along with the “-bnni” option to reduce the risk of overestimating branch support resulting from model violations. We also used a non-parametric bootstrap (BP) with 100 replications to assess branch support and generate BP gene trees for coalescent analyses below. All DNA alignments, protein alignments, and individual gene trees were deposited in Zenodo (10.5281/zenodo.11643025).

### Quantifying phylogenetic properties of individual genes

Each gene was quantified for the following phylogenetic properties using IQ-TREE log files and SortaDate (Smith et al. 2018): number of species, number of parsimony informative sites, average gene tree support, total tree length, root-to-tip branch-length variance, and gene tree–species tree congruence (Table S2). A species tree estimated by ASTRAL-IV v1.19.4.5 (Zhang and Mirarab 2022) based on all 2,135 orthogroups (analysis 15 in Table1) was used for assessing gene tree–species tree congruence, but we removed all Rafflesiaceae and Apodanthaceae species to avoid the systematic bias stemming from artificially fixing this unstable node. Four phylogenetic metrics specific to Rafflesiaceae were extracted: the base compositional bias of Rafflesiaceae, minimum root-to-tip distance among Rafflesiaceae species, the length of the short ascending parent branch of stem-group Rafflesiaceae, and the branch support of Rafflesiaceae. We similarly defined and extracted four Apodanthaceae-specific metrics. The base compositional heterogeneity of Rafflesiaceae and Apodanthaceae was derived from the Chi-square test in IQ-TREE and reported in its log file, whereas the other metrics were summarized using a custom Python script from gene trees (available on GitHub). Finally, we modeled the pairwise Pearson correlation among these phylogenetic metrics. The correlation heatmap was generated using the function “heatmap” from the R package corrplot (Wei et al. 2017).

### Generation of phylogenetic hypotheses with data subsampling

Twenty-two explorative phylogenomic analyses were conducted (Table1, Table S4). These comprised concatenation (10 analyses) and coalescent methods (12 analyses) that used either DNA (18 analyses) or protein alignments (4 analyses) for the complete dataset (17 analyses) or subsampled datasets (5 analyses). Two data subsets were created in an attempt to select an a posteriori set of phylogenetic informative loci and to select clock-like genes with less heterotachy to apply computationally intense mixture models. First, a 446– locus dataset (referred to as G446) was selected from the complete dataset (referred to as G2135) to include loci where Rafflesiaceae and Apodanthaceae formed a monophyletic clade based on ML gene trees estimated with codon-based partition and optimal substitution model determined by ModelFinder (Table S2; Data S1). Second, an 829–locus dataset (referred to as G829) was created to select genes ranked in the top 50% by SortaDate and included more than 32 (71%) species (Table S2; Data S1). A detailed description of the command line used to perform the 22 analyses was provided on GitHub (https://github.com/lmcai/Rafflesiaceae_phylogenomics/tree/master/4_phylogeny_hypo_generation).

Concatenation analysis—We inferred the ML phylogeny based on the DNA and protein alignments with various combinations of partition schemes and substitution models using IQ-TREE. For all IQ-TREE inferences, branch support was evaluated using 1000 UFBP replicates. Analyses 1–4 represented edge-linked partitioned DNA analyses with increasing data filtering effort. The best partition model was determined by the greedy strategy from ModelFinder implemented in IQ-TREE (Kalyaanamoorthy et al. 2017) where partitions were iteratively merged until the model fit did not increase any further (-m MFP+MERGE). Analyses 1 and 2 used DNA alignments filtered by HmmCleaner with a threshold of 50 and a gene-based (Analysis 1) or codon-based (Analysis 2) initial partition. Analysis 3 used the same DNA alignments and a codon-based initial partition, but with the third codon removed. Analysis 4 used DNA alignments without the third codon, filtered by HmmCleaner with a more stringent threshold of 10 and a codon-based partition. Analysis 5 implemented an edge-unlinked mixture GHOST model that can effectively handle heterotachy by assigning sites into individual classes with unlinked model parameters and edge lengths (Lopez et al. 2002; Crotty et al. 2020). We applied this GHOST model with four mixture classes to the DNA alignment filtered by HmmCleaner with a threshold of 50 due to the high RAM usage of this model. The last DNA concatenation (Analysis 6) used a posteriori set of 446 loci (G446) where Rafflesiaceae and Apodanthaceae were monophyletic in individual gene trees. The species tree was then inferred based on a gene-based partition analysis in IQ-TREE.

Analysis 7-10 were protein-based analyses largely reflecting the data filtering strategies of the DNA analyses. Analysis 7 and 8 applied gene-based partition schemes and protein alignments filtered by HmmCleaner with a threshold of 50 and 10, respectively. The site-specific profile mixture model implemented in Analysis 9 was applied to the subsampled G829 dataset and 30 mixture components (LG+C30+F+G) due to the high RAM usage of this model. This PMSF model approximates the empirical profile mixture model of Le et al. (2008), which is a variant of the infinite mixture model CAT-GTR in PhyloBayes. It can effectively ameliorate long-branch attraction artifacts and is computationally more efficient (Wang et al. 2018). Finally, Analysis 10 applied the gene-based partition to the G446 dataset where Rafflesiaceae and Apodanthaceae were monophyletic.

Coalescent analysis—We used ASTRAL-IV and MP-EST v3.0 (Liu et al. 2010) to infer species trees under the coalescent model. For both ASTRAL and MP-EST analyses, ML gene trees of the G2135 dataset inferred from the five sets of partition schemes and substitution models described above were used. Branch support was evaluated using the local posterior probability in ASTRAL and 100 non-parametric bootstrapping in MP-EST. In addition to the complete dataset, we also inferred species trees using the G446 dataset in both ASTRAL and MP-EST.

### Evaluating gene and site phylogenetic conflicts

We used gene and site concordance factors to quantify phylogenetic conflicts using IQ-TREE. The gene concordance factor (gCF) is defined as the percentage of decisive gene trees containing that branch (Minh, Hahn, et al. 2020). The site concordance factor (sCF) was similarly defined but used a heuristic method to sample alignment sites while minimizing the effects of homoplasy and taxon sampling (Mo et al. 2023). To calculate gCF and sCF in IQ-TREE, we used the species tree, gene trees, and DNA alignments from the coalescent-based Analysis 15 (Table 1). These gene trees were inferred under the optimal substitution models determined by ModelFinder without codon-based partition. For sCF calculation, 200 quartets were sampled to obtain stable site concordance factors (--scfl 200) and we inferred sCF for both the concatenated DNA alignment and three codon positions separately.

### Fit of gene trees and sites to proposed topologies

The initial phylogenetic reconstructions revealed several possible placements of Rafflesiaceae and Apodanthaceae associated with Euphorbiaceae+Peraceae, Putranjivaceae, Pandaceae, Erythroxylaceae+Rhizophoraceae, Ixonanthaceae, and Linaceae. We tested the fitness of individual genes and sites to the 8 alternative topologies commonly recovered from our species inference analyses (Table 1) using the methods described in Shen et al. (Shen et al. 2017).

Briefly, we conducted constrained phylogenetic searches enforcing the relationship imposed by each of the 8 hypotheses but no constraints on the rest of the tree. Branch length and the overall species tree topology were optimized using gene-based partition (without merging) and substitution models determined by ModelFinder. Gene and site likelihood scores for each tree were then obtained under the same partition and model parameters from the previous step using the “-wpl” and “-wsl” flags in IQTREE, respectively. The gene-specific rate was obtained from the IQTREE log file (under “Partition-specific rates”) and the site-specific rate was inferred using the “-wsr” option in IQTREE. These gene and site likelihood scores and rates were reported in Data S3.

### Identify spurious relationships arising from fast-evolving genes

To explore how fast-evolving genes may lead to spurious species tree reconstruction, we first ordered the 2,135 loci based on the gene-specific rate for both DNA and protein alignments (Data S3). We then examined the changes in cumulative difference of log likelihoods (ΔLL) between the best-supported (H1 for DNA and H3 for protein) and alternative topologies when more fast-evolving genes were added to the concatenated matrix (Fig. 3). Results robust to evolutionary rates should demonstrate a steady increase in ΔLL (e.g., Fig. 3C). In contrast, results sensitive to evolutionary rates are expected to have ΔLL curves with less predictable trends and even sharp changes when the fastest-evolving genes are added (e.g., Fig. 3B). To further evaluate the statistical significance of these deviations, we generate 1000 bootstrap replicates where genes were randomly ordered and plotted the same ΔLL curve. If the empirical curve falls out of the 95% confidence interval, we consider a significant influence of rates to generate potentially spurious relationships.

### Testing the anomaly zone

The anomaly zone in a four-taxon species tree is defined by the length ratio of the two internal branches (Degnan and Rosenberg 2006). To test if the focal clades fall within the anomaly zone, we estimated the boundaries of the anomaly zone a(x) according to equation 4 from (Degnan and Rosenberg 2006). We used the phylogeny estimated by ASTRAL from the complete dataset as the reference species tree (Analysis 15 in Table 1). Here, the three internal nodes among the focal clades form three pairs of parent-descendent branches whose topology aligns with the asymmetrical four-taxon phylogeny from (Degnan and Rosenberg 2006). If the length of the descendant internal branch is smaller than a(x), then the internode pair is in the anomaly zone.

### Coalescent simulation

To further explore how ILS and homoplasy affect species tree reconstruction, we simulated gene trees and DNA alignments based on the coalescent and mutational parameters from the empirical dataset. The MP-EST species tree estimated from the complete dataset (analysis 14 in Fig. 2) was used as the reference species tree and was pruned to include only one species from each of the focal clades plus a non-Malpighiales outgroup, Crossopetalum. We used the function “sim.coal.mpest” in the R package Phybase v2.0 (Liu and Yu 2010) to simulate 10,000 gene trees using the reference species tree. We then modified the branch lengths of the simulated gene trees to reflect the inter-species and inter-locus rate variation in mutational units. To do so, we fit a Gaussian distribution to each internal and terminal branch (e.g., terminal branches leading to Sapria, Rafflesiaceae) from the empirical ML gene trees using function “fitdistr” in the R package MASS v7.3-54 (Ripley et al. 2013). We then assigned branch length in mutation units to the simulated trees by randomly sampling the corresponding Gaussian distribution. Finally, we used the parameters of the GTR+F+G4 model estimated by IQ-TREE based on the complete dataset (analysis 1 in Fig. 2) to simulate DNA alignments in Seq-Gen v1.3.4 (Rambaut and Grassly 1997). For each gene, four alignments with different lengths (500, 1000, 1500, and 2000 bp) were simulated to reflect high to low levels of gene tree estimation error (Mirarab et al. 2016; Cai et al. 2021b). We then inferred gene trees in IQ-TREE under the GTR+F+G4 model using 100 bootstrap replicates. Species trees were estimated using both concatenation and coalescent methods. For concatenation analyses, we applied the unpartitioned dataset in IQ-TREE under the GTR+F+G4 model. Branch support was estimated by 1000 UFBP. For the coalescent method, we applied the ML gene trees in MP-EST and ASTRAL. Branch support was estimated by 100 BP replicates. We also subsampled the dataset to include 500, 1000, 2000, 3000, 4000, and 5000 loci with four alignment lengths. Finally, we conducted AU tests in IQ-TREE and likelihood ratio tests in MP-EST to statistically evaluate alternative topologies.

### Divergence time estimation

To fully utilize our phylogenomic data for divergence time estimation, we applied both the penalized likelihood method implemented in treePL v1.0 (Smith & O’Meara, 2012) and the Bayesian method implemented in PhyloBayes v4.1 (Lartillot et al. 2009). For both analyses, we fixed the topology using the species tree derived from the ASTRAL analysis 15 (Table 1).

For the treePL analysis, branch lengths of the species tree were inferred using the concatenated DNA alignments, partition scheme, and substitution model from analysis 5 (Table 1) in IQ-TREE. This input alignment included all 2,135 loci, masked using HmmCleaner with a threshold of 10, and excluded all third codon positions. We fixed the root age at 118 Ma to reflect the crown group age of the Celastrales-Oxalidales- Malpighiales clade from Ramírez-Barahona et al. (Ramirez-Barahona et al. 2020). In addition, eight fossils in Malpighiales and one fossil from Oxalidales were used to constrain the minimum age of the clade they each represented (Table S8). We then primed and cross-validated our data to establish the best smoothing parameter (smooth = 0.01). Finally, we conducted a thorough search to infer the best time tree.

For the PhyloBayes analysis, we similarly applied the concatenated DNA alignment without the third codon position and masked with HmmCleaner using a threshold of 10. To ease with convergence, we used the clock-like G829 loci. Divergence time estimation was conducted under the log-normal autocorrelated relaxed clock model and the CAT Dirichlet process mixture for among-site substitution heterogeneities. We constrained the root age using a uniform distribution with a lower and upper bound of 89 and 125 Ma, respectively. These boundaries were determined by the age of the oldest fossil Palaeoclusia chevalieri (89 Ma) (Crepet and Nixon 1998) within Malpighiales and the oldest angiosperm macrofossil Archaefructus liaoningensis at 125 Ma (Sun et al. 1998). For internal node age calibrations, we used a uniform distribution for each fossil (Table S8) where the fossil’s age served as the minimum and 125 Ma as the maximum. We ran two Monte Carlo Markov chains per model and kept every tenth sample until 30,000 samples were collected.

Convergence was checked by computing the discrepancies and effective sample sizes of the posterior averages and visualizing the trend of the posterior. Consensus trees were computed by removing 30% of the burn-in trees.

## Acknowledgements

We thank the two anonymous reviewers for their insightful comments on the initial version of the manuscript. We thank Steven R Manchester, Peter Wilf, and members of the Davis lab for feedback on early versions of this study.

## Data availability

All DNA alignments, protein alignments, and individual gene trees were available on Zenodo (10.5281/zenodo.11643025). All codes to conduct the analysis were available on GitHub https://github.com/lmcai/Rafflesiaceae_phylogenomics.

