## Supplementary Information for "The danger zone: the joint trap of incomplete lineage sorting and long-branch attraction in resolving the Gondwanan origin of Rafflesiaceae and Apodanthaceae"

**Supplementary figure**

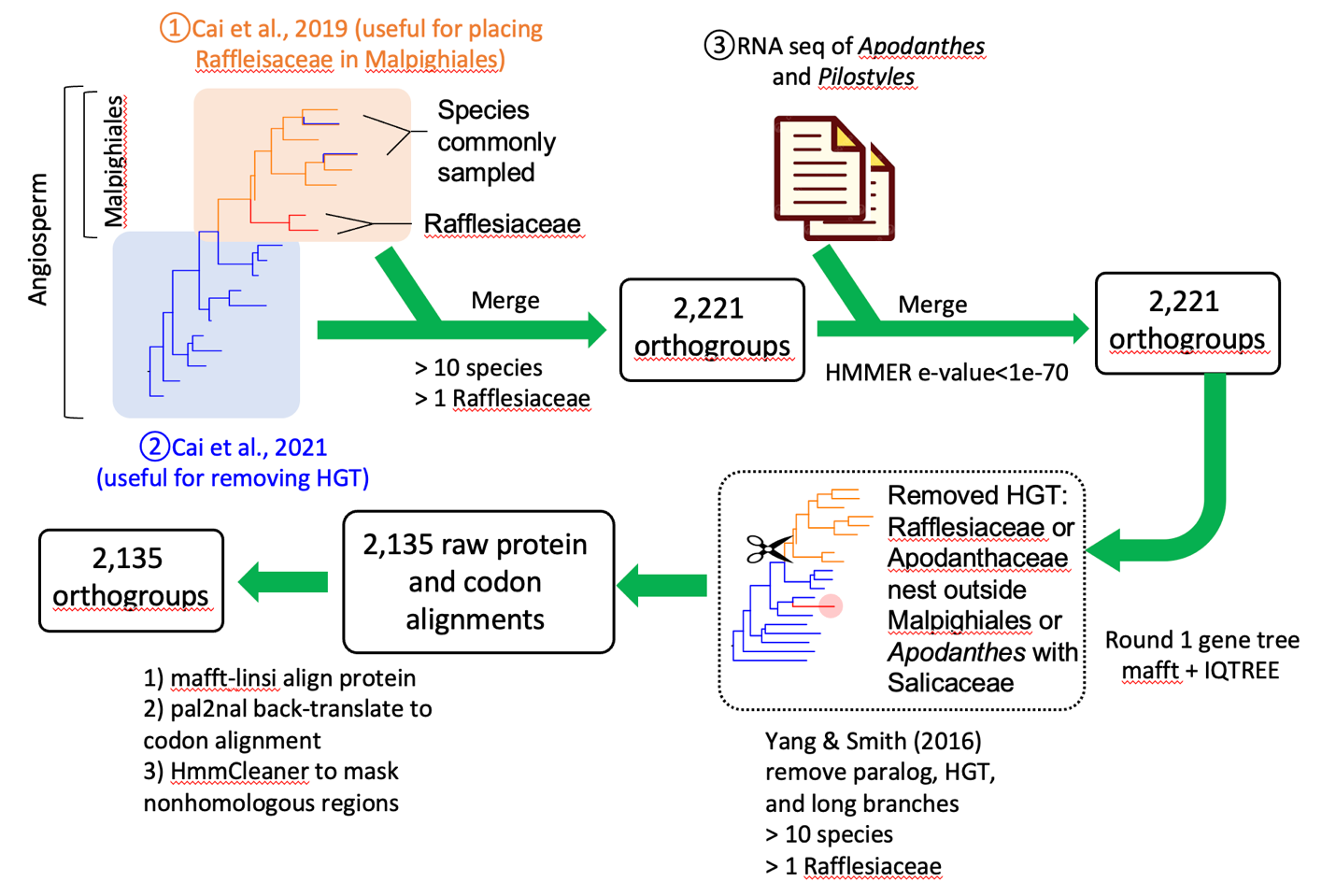

**Fig S1** Bioinformatic pipeline for phylogenomic dataset assembly. Three sets of data, including the newly published Apodanthaceae RNA sequences, were merged into orthogroups based on sequence similarity. A phylogeny-based method from Yang and Smith (2016) was subsequently applied to remove paralogs and horizontal gene transfers. The final codon alignments were cleaned using HmmCleaner to mask non-homologous regions.

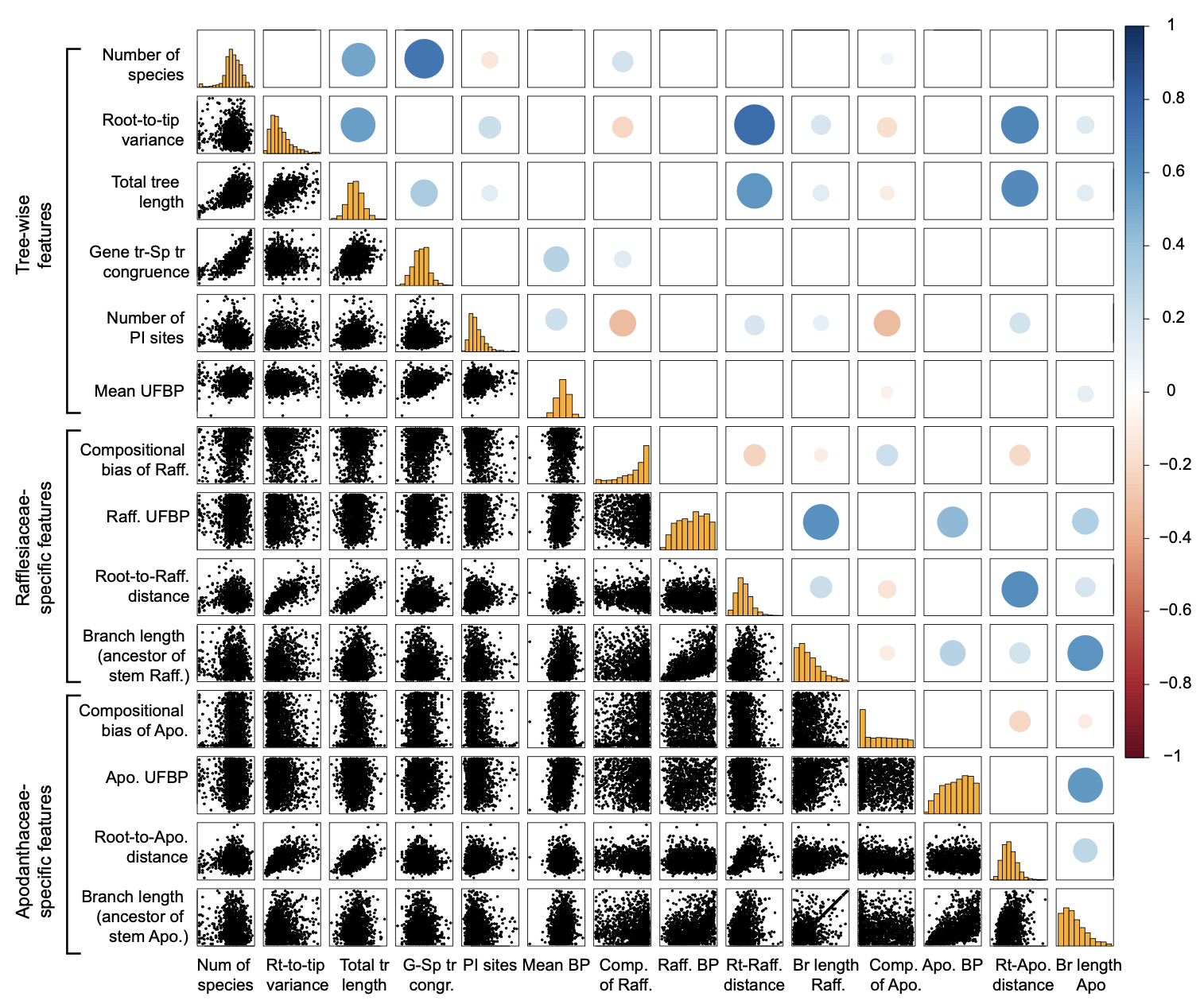

**Fig S2** Pairwise correlation between fourteen metrics of gene phylogenetic property. The scatter plots on the lower left depict the raw pairwise distribution; the circles on the upper right represent the significance and direction of Pearson correlation tests. The size of the circles is negatively scaled with the p-value, and the red-to-blue hue of the Pearson correlation coefficient represents the negative-to-positive correlation.

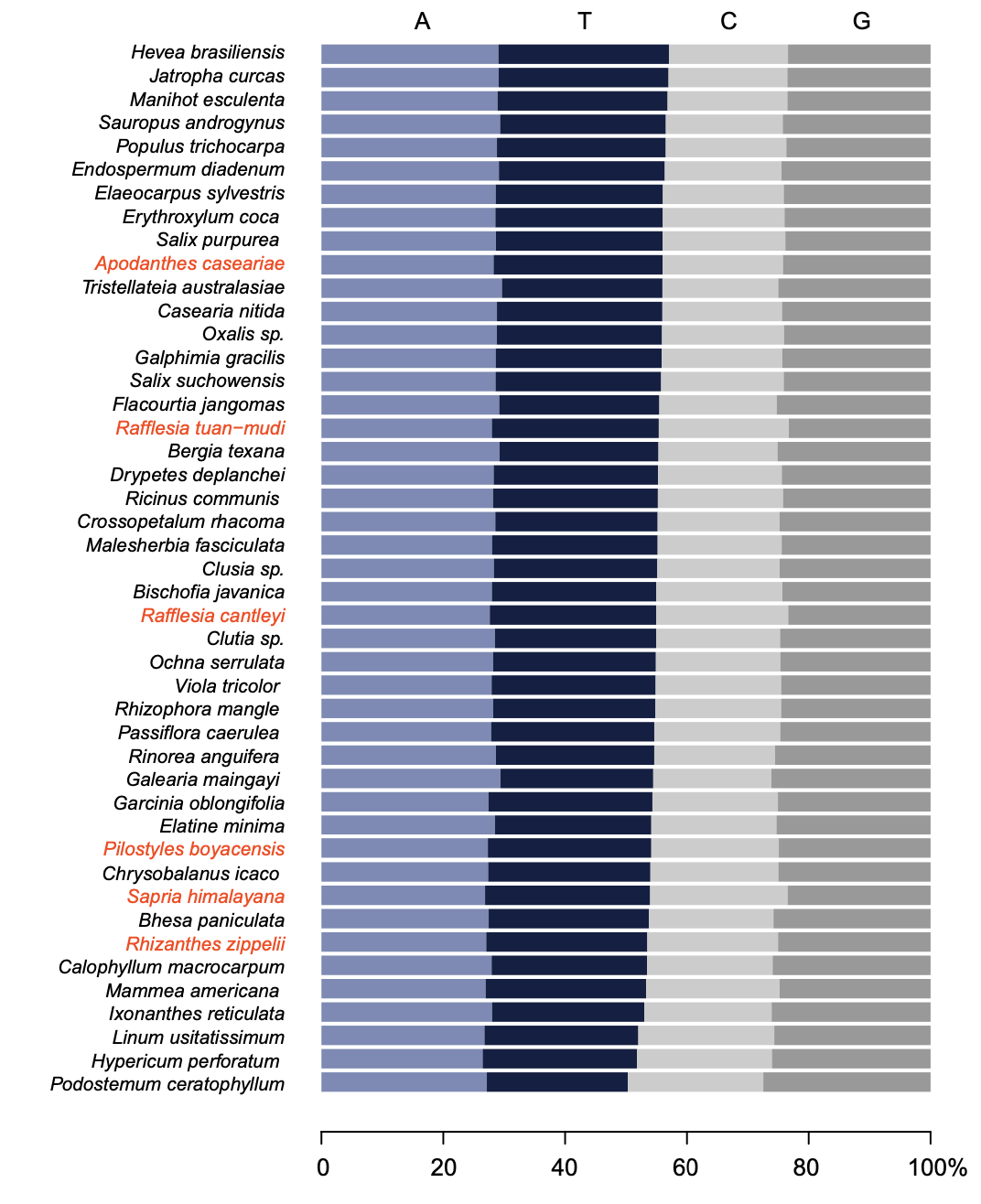

**Fig. S3** DNA composition of Malpighiales species summarized from 2135 genes. Species are ranked based on AT% from high to low. Rafflesiaceae and Apodanthaceae species are highlighted in red.

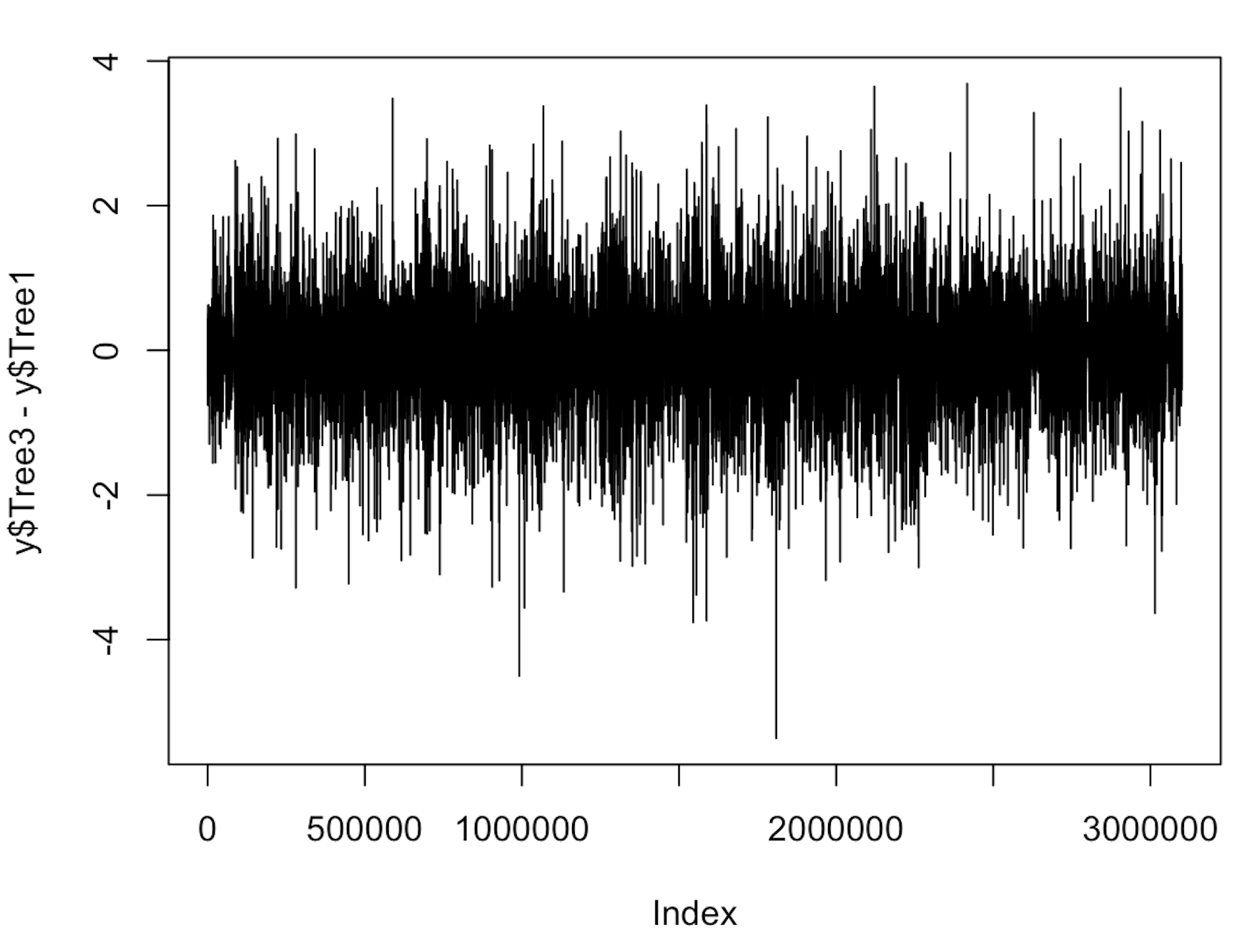

ΔLL (H1-H3)

**Fig S4** One example of the difference in log likelihoods (ΔLL) between alternative topologies across all 3 million DNA sites in the complete dataset G2135. The plot shows the ΔLL between H1 and H3. The site-specific likelihoods were inferred using the ‘-wsl’ flag in IQ-TREE.

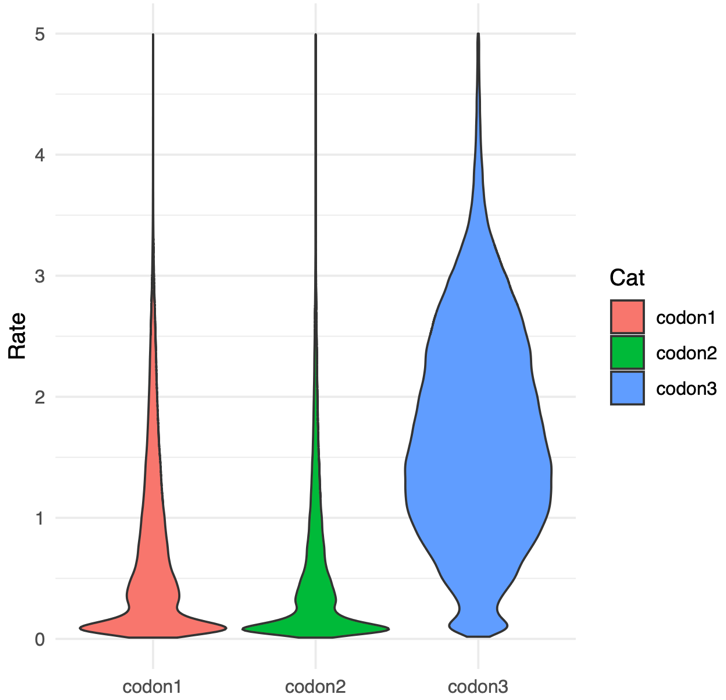

**Fig S5** Distribution of site rate for three codon positions in the complete dataset. The site rates were inferred using the ‘-wsr’ flag in IQ-TREE. The third codon has significantly higher rates compared to the first and second codons.

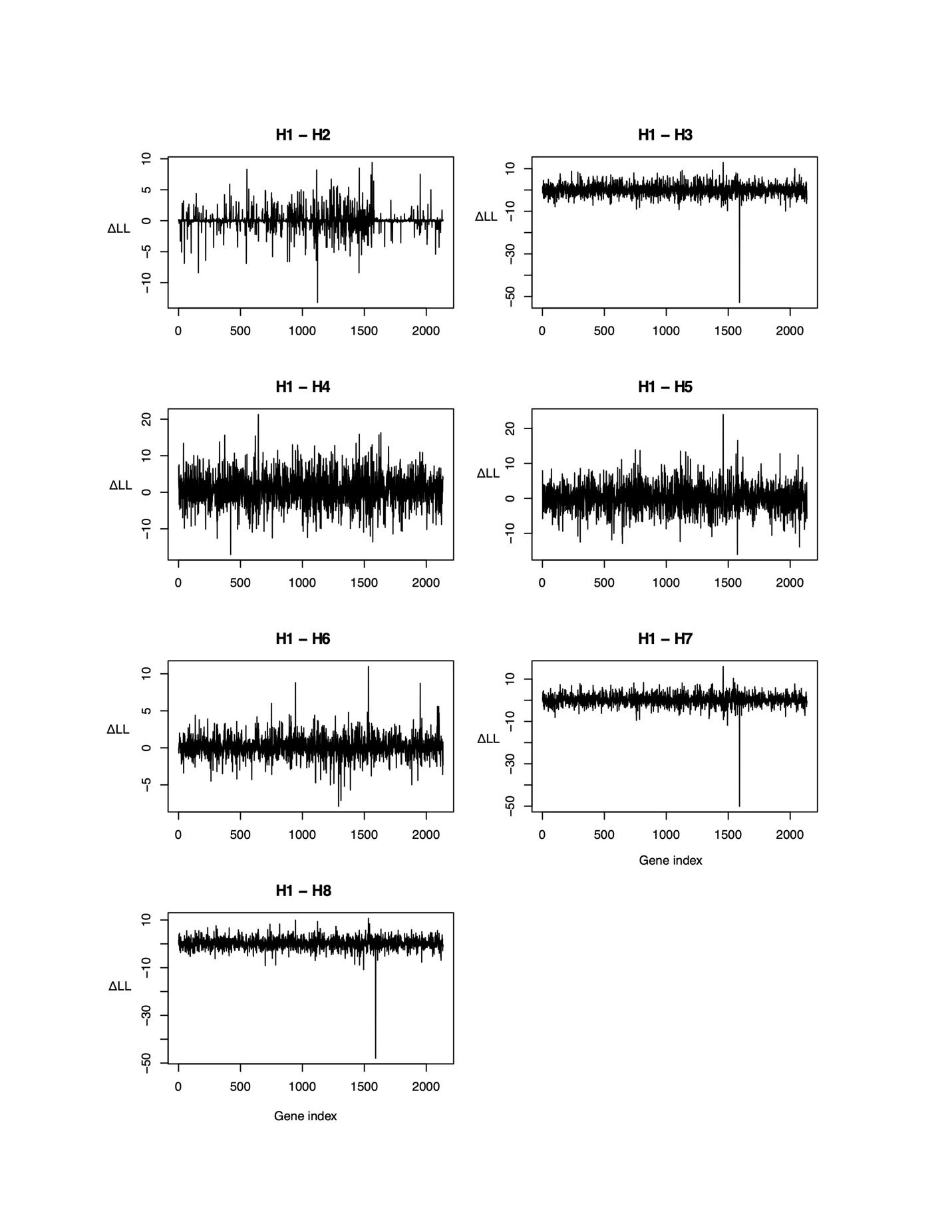

**Fig S6** Difference in log likelihoods ΔLL between the best-supported tree H1 and the other seven topologies across all 2,135 loci based on DNA alignments. Note that “Locus 313” has exceptional ΔLL values when comparing H1 to H3, H7, and H8. These locus-specific likelihoods were inferred using the ‘-wsl’ flag in IQ-TREE.

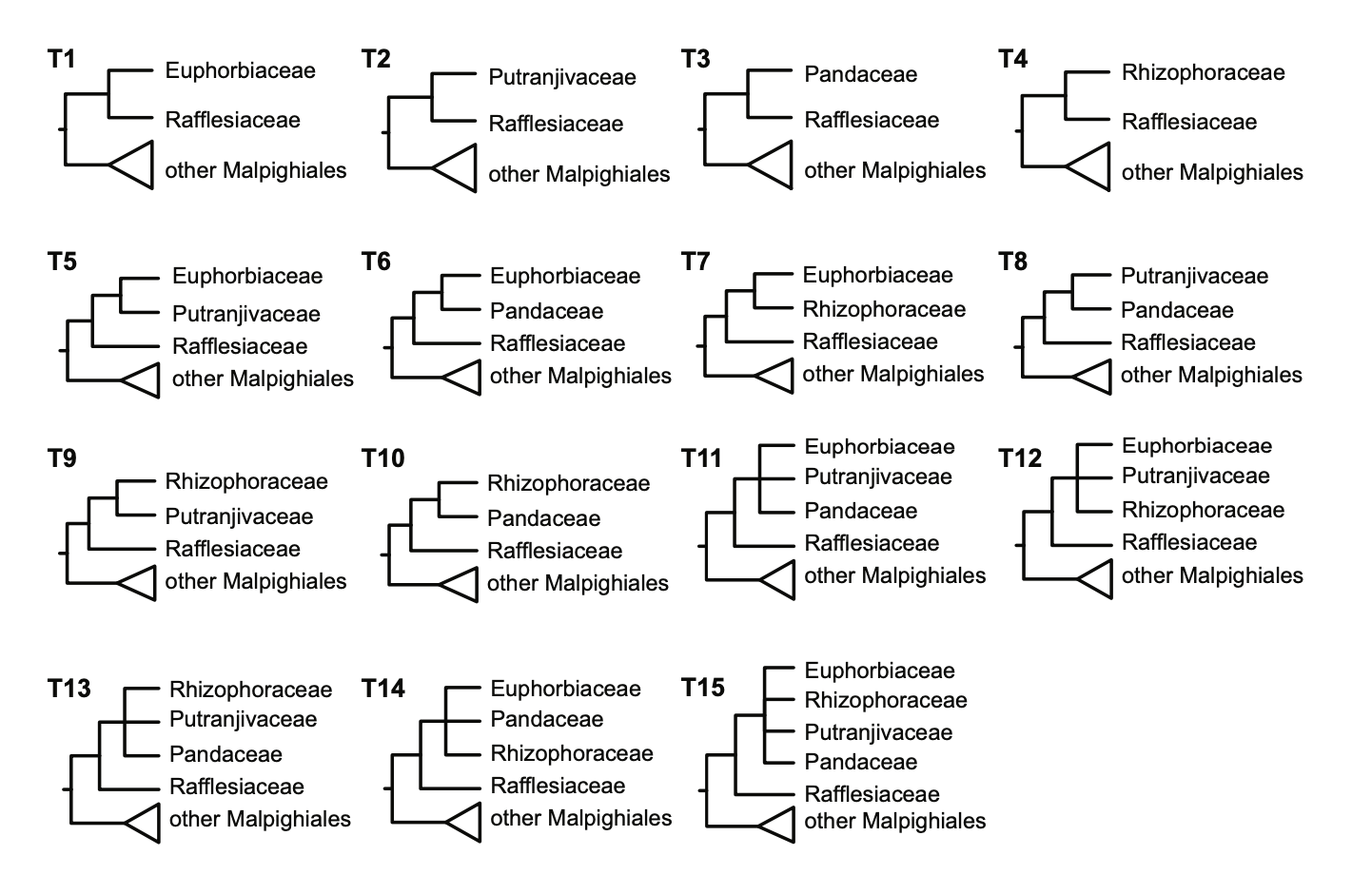

**Fig S7** Topology for all 15 possible placements of Rafflesiaceae within a simulation experiment of five species.

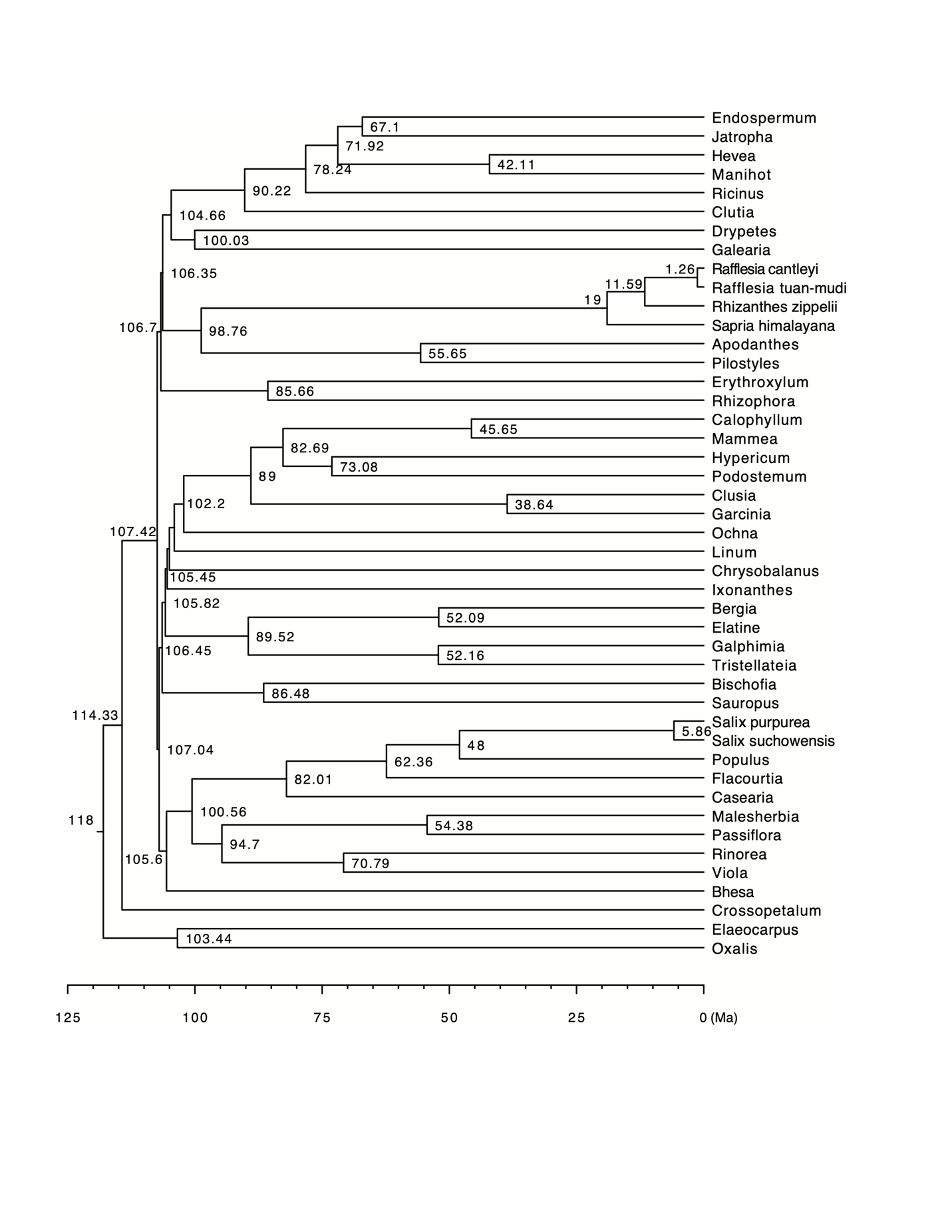

**Fig S8** Divergence time of Malpighiales inferred from the penalized likelihood method implemented in TreePL. The time tree is inferred using the concatenated DNA matrix of the complete dataset with the third codon removed and nine fossil calibration points. The numbers on the node indicate divergence time in millions of years ago (Ma).

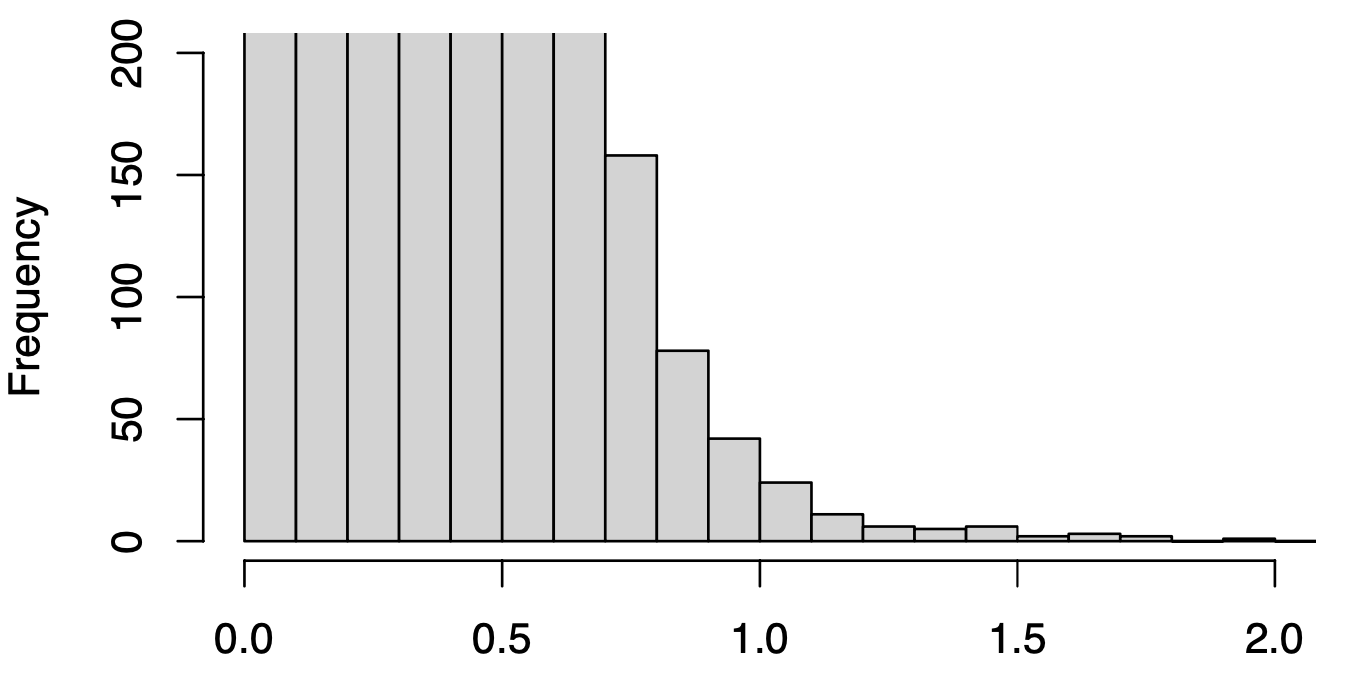

**Fig S9** Distribution of branch lengths across 2,135 loci. Based on this distribution, we pruned branches longer than 1.5 as a sequence filtering step for data cleaning. The top part of the histogram is truncated to facilitate visualization.

**Supplementary Table**

**Table S1** Taxon sampling and source of sequence data. The species name, taxonomic affiliation, and data source are provided for 45 species included in the current study.

**Table S2** Phylogenetic properties of 2135 genes. These metrics include nine tree-wise features for each gene, including the number of species, alignment length, number of phylogenetic informative sites, number of phylogenetic invariable sites, average branch support measured by ultrafast bootstrap values (UFBP) in IQTREE, median root-to-tip distance, total tree length (measured by SortaDate), root-to-tip branch length variance (measured by SortaDate), and gene tree–species tree congruence (measured by SortaDate); four metrics measured specifically for Rafflesiaceae and Apodanthaceae respectively: DNA compositional bias (chi-square test *p*-values), the length and support of the stem group, and minimum root-to-tip distance.

**Table S3** List of genes where Rafflesiaceae and Apodanthaceae are monophyletic in gene trees. “Y” means Rafflesiaceae and Apodanthaceae are both present and monophyletic; “N” means Rafflesiaceae and Apodanthaceae are both present but not monophyletic; “NA” means missing one of Rafflesiaceae and Apodanthaceae and cannot evaluate. Each gene alignment was analyzed using five substitution models: (1) GTR_codon: codon-based partition and forcing the GTR substitution model *without* rate heterogeneity across sites (i.e., no +I or +R or +G); (2) msetGTR_codon: codon-based partition and forcing the GTR substitution model *with* rate heterogeneity across sites; (3) MFP_codon: codon-based partition and using the optimal substitution models determined by ModelFinder; (4) msetGTR: no partition within loci and forcing the GTR substitution model with rate heterogeneity across sites; (5)MFP: no partition within loci and using the optimal substitution models determined by ModelFinder.

**Table S4** Phylogenetic analysis schemes for 22 concatenation and multispecies coalescent model based methods. Columns show the input data type, number of genes, evolutionary models, and phylogenetic placement of Rafflesiaceae+Apodanthaceae for each analysis.

**Table S5** Topology tests for alternative placements of Rafflesiaceae using various data subsets based on the Approximately Unbiased (AU) test and bootstrap proportion using RELL method (bp-RELL). Hypotheses 1 to 8 (H1-H8) are defined in Fig. 1 and the log likelihood (LL) is inferred from IQTREE using the same best partitioning scheme for all eight topologies. Topologies rejected by the corresponding statistical test are marked by “–”.

**Table S6** Topology test on simulated data using concatenated DNA alignments. Hypotheses 1 to 15 (H1-H15) are defined in Fig 1 and H11 is used as the true species tree for simulation. AU-test *p-*values are calculated using concatenated DNA matrices of different sizes (a-d). The lengths of simulated genes vary from 500 bp (a), 1000 bp (b), 1500 bp (c), to 2000 bp (d).

**Table S7** Topology test on simulated data using the coalescent based likelihood ratio test. Hypotheses 1 to 15 (H1-H15) are defined in Fig 1 and H11 is used as the true species tree for simulation. Differences in pseudo-likelihoods between the inferred best tree and the true species tree (H11) are calculated for datasets with different gene lengths and numbers (a-d). For the biggest dataset (gene length =2000 bp; d), we calculate the likelihood ratio test *p-*values for a randomly selected alternative tree H5, which cannot be rejected in any test.

**Table S8** Fossil calibration points for divergence time estimation. The clade definition, age, and reference are listed for 9 fossils used to calibrate the time tree of Malpighiales.

**Supplementary Data**

**Reviewer link:**

[**https://zenodo.org/records/15587049?token=eyJhbGciOiJIUzUxMiJ9.eyJpZCI6ImMyMWYwMTU0LTgyMzktNDA2MS05NzVkLTBkYTI4YTBhZmIzOCIsImRhdGEiOnt9LCJyYW5kb20iOiJmNmIyMjQwNmIyZDdiYmQzZWM3YWFkMjI1YzU2M2QwYiJ9.0PiyXOGRt3kmJ_CbyOv4A0X8W0IHWqFuxSzHIzE5COp33hOF6dQJ3UyPQMrJWjjuaqaVrFNFSLotSKQlj4vE4Q**](https://zenodo.org/records/15587049?token=eyJhbGciOiJIUzUxMiJ9.eyJpZCI6ImMyMWYwMTU0LTgyMzktNDA2MS05NzVkLTBkYTI4YTBhZmIzOCIsImRhdGEiOnt9LCJyYW5kb20iOiJmNmIyMjQwNmIyZDdiYmQzZWM3YWFkMjI1YzU2M2QwYiJ9.0PiyXOGRt3kmJ_CbyOv4A0X8W0IHWqFuxSzHIzE5COp33hOF6dQJ3UyPQMrJWjjuaqaVrFNFSLotSKQlj4vE4Q)

**Data S1** Sequence alignments and gene trees for the 2135 loci analyzed in the study. The lists of loci included in the data subsets G446 and G829 are also provided. Both the raw alignments and masked alignments by HmmCleaner are provided for DNA and protein sequences. Each gene has 5 sets of gene trees inferred using different models in IQTREE:

(1) *.GTR_codon.treefile: codon-based partition within gene (first, second, third codon in separate partitions) + forcing GTR (no +I or +R or +G); (2) *.MFP_codon.treefile: codon-based partition within gene + ModelFinder best fitting model; (3) *.msetGTR.treefile: no partition within gene + forcing GTR (with +I or +R or +G); (4) *.msetGTR_codon.treefile: codon-based partition within gene + forcing GTR (with +I or +R or +G); (5) *.na.mask.fas.treefile: no partition within gene + ModelFinder best fitting model.

**Data S2** Species tree in newick format generated by the 22 analyses described in Cai et al. Table 1 and Table S3. For partition-based concatenation analyses, the best partition and substitution scheme in nexus format from IQ-TREE is also provided.

The program and command line used to generate these species trees can be found on GitHub: https://github.com/lmcai/Rafflesiaceae_phylogenomics/tree/master/4_phylogeny_hypo_generation

**Data S3** Log Likelihood per gene, log Likelihood per site, and rate per site estimated from the concatenated DNA sequence of the complete dataset. These likelihoods and rates are used for AU tests in Table S5 and the rate-sensitivity analysis in Fig. 3.

**Supplementary Note**

**Supplementary Note 1**

Phylogenomic dataset assembly

We obtained the coding sequences, protein sequences, and gene trees from 5,112 orthogroups used in the phylogenomic study of Malpighiales (Cai et al. 2019) and 6,552 orthogroups used in the genome research of *Sapria himalayana* (Cai et al. 2021). The former study sampled 36 Malpighiales species and three outgroups (Table S1) to investigate whole genome duplications within Malpighiales (Cai et al. 2019). The latter study sampled four Rafflesiaceae, three free-living Malpighiales, and 31 broadly-sampled angiosperm species to identify HGT within Rafflesiaceae (Cai et al. 2021). Based on the Ensembl gene IDs of the three free-living Malpighiales species commonly sampled in both studies (*Manihot esculenta* Crantz, *Populus trichocarpa* Torr. & A.Gray ex Hook., and *Jatropha curcas* L., Table S1), we first merged the sequence alignments to generate a combined phylogenomic dataset containing 43 species. This yielded 3,086 orthogroups for further examination.

To add the two Apodanthaceae species into the dataset, we first generated preliminary DNA alignments for each of the 3,086 orthogroups using MAFFT v.7.299 (Katoh and Standley 2013). These preliminary alignments were used to build profile hidden Markov models to search for orthologous sequences within *Apodanthes* and *Pilostyles* using HMMER v3.4 (Mistry et al. 2013). We then identified the best HMMER hit with e-values lower than 1e-70 as potential orthologs.

After merging sequences from all three studies, we performed DNA alignment for 3,086 orthogroups in MAFFT. A preliminary gene tree was also inferred for each orthogroup using the default setting in IQ-TREE (iqtree2 -s). We applied a tree-based approach to select vertically inherited Rafflesiaceae and Apodanthaceae genes nested within Malpighiales using the python script ‘prune_paralogs_RT.py’ from Yang and Smith (2014). This step removes paralogs and horizontal gene transfers from non-Malpighiales lineages such as Vitaceae and Fabaceae, which are the hosts of Rafflesiaceae and *Pilostyles*. We also used a custom Python script to remove *Apodanthes* sequences clustered with their host Salicaceae within Malpighiales. In addition, we removed exceptionally long branches with root-to-tip distances larger than 1.5, the threshold of which is determined by the distribution of branch lengths (Fig. S9).

We filtered the 3,086 orthogroups to include at least one vertically transmitted gene from Rafflesiaceae or Apodanthaceae and at least 10 species (23% species completeness) in total. This yielded 2,135 orthogroups. We realigned the protein sequences of these 2,135 loci using the MAFFT-ensi algorithm (mafft --genafpair --maxiterate 1000) and then converted them into the corresponding codon alignments using pal2nal (Suyama et al. 2006). Finally, protein and DNA alignments were masked using the profile hidden Markov models implemented in HmmCleaner v0.180750 (Di Franco et al. 2019). We applied a threshold of 50 and 10 to generate two sets of alignments with relaxed or stringent site filtering, respectively.
